# Unravelling neuromechanical constraints to finger independence

**DOI:** 10.1101/2025.05.23.655831

**Authors:** Daanish M. Mulla, Paul M. Tilley, Peter J. Keir

## Abstract

Intentional use of a single finger results in involuntary forces and movements among other fingers. Constraints to finger independence are attributed to both neural and mechanical factors, but the contribution of these factors is debated. We hypothesized that neural factors primarily constrain finger independence during isometric exertions whereas mechanical factors impose larger constraints during movements. We investigated changes in finger independence following a ring finger fatigue protocol. We assumed that with fatigue, the ability to actively transmit forces across fingers through neural pathways will be reduced but force transmission passively through mechanical pathways will remain unaffected. Participants performed isometric finger contractions and flexion-extension movements at baseline and following a ring finger fatigue protocol. At baseline, involuntary ring finger forces ranged from 7.3-16.5% MVC. Consistent with our predictions, involuntary ring finger forces decreased by 2.5-8.9% MVC following fatigue. In contrast, involuntary ring finger movement did not change or surprisingly in several cases, increased by greater than 10-20° following fatigue relative to baseline across movement tasks. Our findings demonstrate that the neuromechanical control of finger force versus motion are distinct from each other and can alter the constraints to finger independence in a task-dependent way.

## 1. Introduction

Humans can perform a rich repertoire of tasks using their hands, ranging from activities involving delicate finger control to forceful grasps. Despite dexterous manipulative abilities, we do not possess complete ability to independently control each of our fingers. When individuals are instructed to exert force with or move a single finger, involuntary finger forces [1–5] and movements [6, 7] are observed across non-task fingers.

The constraints to finger independence are attributed to mechanical and neural factors [8–10]. From a mechanical standpoint, the musculotendon units of different fingers are physically coupled through connective tissue linkages. On the extensor side, these connections include the prominent juncturae tendineii between the tendons of the extensor digitorum communis (EDC) [11–13]. On the flexor side, inter-tendinous connections can be found between the flexor digitorum profundus (FDP) [14–16] and superficialis (FDS) [17] of the different fingers.

Additionally, common myofascial sheaths, shared neurovascular tracts, and fused aponeuroses between extrinsic finger muscle bellies [11, 18–21] as well as the finger web spaces [13] are additional sources for passive force transfers. Single finger forces and movements will lead to relative changes in musculotendon length and position between the task finger and non-task fingers, consequently straining connective tissue linkages and transmitting passive forces onto non-task fingers [20, 22]. Direct evidence for these passive force transfers is exemplified by the abolishment of non-task finger movements with excision of connective tissue linkages within cadaveric specimens [13, 16]. From a neural standpoint, hard-wired pathways within the central and peripheral nervous system may drive common synaptic input to the different compartments of the FDS, FDP, and EDC, leading to an inability to independent activate the muscles of a single digit. Neurons in the primary motor cortex can be tuned to multiple fingers, with extensive overlap in the spatial activations during single finger tasks [23, 24]. Further, last order inputs from a single neuron to the spinal cord can diverge onto multiple motoneuron pools [10, 25]. Peripherally, a motoneuron pool can synapse onto multiple muscles [26–28]. In support of these neural pathways, the different compartments of the FDP, FDS, and EDC muscles exhibit patterns of short-term synchronous motor unit firing [29–31].

The contribution of mechanical and neural constraints to finger independence is heavily debated and complicated by conflicting findings in the literature studying different task conditions. Some scientists argue that the role of mechanical constraints is relatively minor, with two often cited studies by Kilbreath & Gandevia (1994) [14] and Keen & Fuglevand (2003) [32]. In the former study [14], passive rotation of the distal interphalangeal (DIP) joint in anesthetized hands did not produce movements of non-task fingers, suggesting limited passive force transfers between the FDP compartments. It is important to note that DIP rotation causes relatively small displacements of the FDP tendon (∼6 mm for 80° joint angle rotation), which is approximately half the magnitude when compared to either proximal interphalangeal (PIP) or metacarpophalangeal (MCP) joint rotations [33]. The limited tendon displacement with isolated DIP joint rotation may not sufficiently strain connective tissue linkages between FDP compartments to passively transmit any forces across fingers. Supporting this argument, passive single finger flexion-extension cycles of the MCP joint produces movements of non-task fingers, which were not significantly different compared to active MCP rotations [7]. Further, passive single flexion of the whole finger (MCP, PIP, and DIP joints) generates flexion forces across non-task fingers when held statically in cadaveric hands [34]. In the second often cited study as evidence for minor contributions of mechanical constraints [32], electrical stimulation of the EDC primarily transmitted forces to a single finger during resting and low-level (∼10% maximal voluntary force) static contractions. Similar to our reasoning above, as the fingers were fixed isometrically and electrical stimulation generated low magnitudes of finger force (averaged ∼120 mN external force), it is possible that the relative change in length between adjacent EDC tendons was not sufficient for straining connective tissue linkages. In fact, fingers can independently move over small ranges of motion [35], beyond which, greater relative movement between fingers increases non-task finger forces [34, 36]. Altogether, these results suggest that mechanical constraints to finger independence may play a minor role when relative changes in musculotendon lengths and position between fingers may be minimal (e.g., single finger DIP rotation, low-level static contractions) but could grow in importance during tasks eliciting greater strain among connective tissue linkages (e.g., whole single finger movements) [35–37]. To investigate further, we need to study both static and movement tasks within a single experimental paradigm and evaluate whether finger independence can be differentially affected across task conditions.

The overarching purpose of our work is to determine the contribution of neural versus mechanical factors in limiting finger independence. We hypothesized that neural factors are the primary constraints to finger independence during static, isometric contractions whereas mechanical factors impose larger constraints during finger movements. To study our hypothesis, we investigated changes in finger independence following a fatigue protocol targeting the ring finger. Fatigue of the ring finger was targeted because it is the least independent finger (i.e., largest magnitude of involuntary forces and movements) and is expected to be most sensitive to any fatigue-induced changes in finger independence. As fatigue will diminish a muscle’s force generating capacity, we assumed that the ability to transmit forces through neural pathways will be reduced (i.e., same level of activation will lead to reduced force output) but that force transmission passively will remain unaffected. Thus, we predicted that ring finger fatigue will reduce involuntary ring finger forces during instructed isometric exertions of adjacent fingers (middle and little) but will not alter involuntary ring finger movement during instructed movements of adjacent fingers (middle and little).

## 2. Methods

### 2.1. Participant Information

Twenty right-hand dominant participants (10M/10F; age 21.7±3.4 years; height 170.7±7.3 cm; mass 69.3±11.1 kg) from the university population with no self-reported history of upper limb pain and injury were recruited for this study. The McMaster Research Ethics Board approved the study. All participants provided written informed consent prior to participating.

### 2.2. Equipment and Instrumentation

Participants were seated upright with their right arm placed in a custom-designed apparatus affixed to a height-adjustable table (Figure 1a). The height of the table was set such that participants arms were slightly abducted with the elbow flexed at 90°. The apparatus was equipped with wrist and elbow braces to maintain a mid-prone position (i.e., thumb pointing up) with a neutral wrist flexion-extension and radial-ulnar deviation posture (i.e., straight wrist). The apparatus was further equipped with a plate secured with four uniaxial force transducers (MLP50, Transducer Techniques, CA, USA). Each transducer was attached to a finger ring fitted to the participant’s middle phalanx of the index, middle, ring, and little fingers (Figure 1a). The finger rings were 3D printed (polylactic acid) in different sizes to accommodate for participant anthropometrics. The apparatus was only used during isometric contractions (see experimental protocol below), with participants arms removed from the apparatus during movement trials.

**Figure 1:**
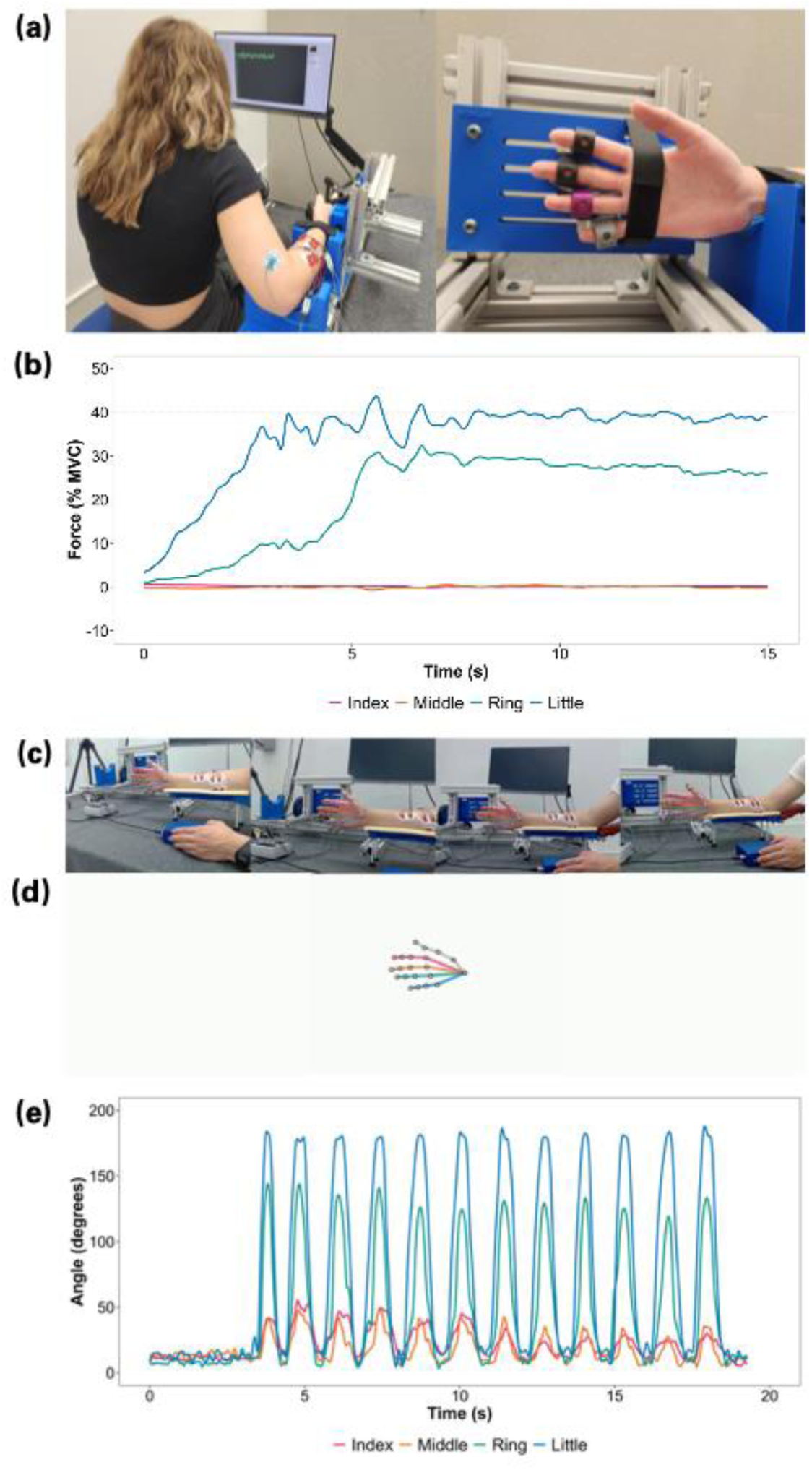
Experimental setup depicting participant positioning for (a) isometric force and (c) movement tasks. (a) Participant was seated with their right arm braced with custom-designed wrist and elbow supports adjusted to individual anthropometrics. Instructed finger force feedback was provided on a monitor placed in front of the participants. Fingers were secured with fitted rings around the middle phalanx attached to force transducers. (b) Finger forces (% MVC) from a sample submaximal isometric task – little finger flexion at 40% MVC. (c) Camera views of the participant performing finger movements with their right hand, with overlaid predictions of 21 hand key points and (d) triangulated 3D positions. (e) Finger joint angles (summed across MCP and PIP) from a sample movement trial – little finger flexion-extension performed at 0.75 Hz.

Finger movements of the right-hand were recorded using four webcams (Logitech C920e, Logitech, Newark, CA, USA) (Figure 1c). Videos were recorded at 480×640 pixels with a frame rate of 30 Hz (vMix, StudioCoast). Camera calibration was performed using a 11×8.5-inch checkerboard (6×8 squares with a checker size of 31.1 mm) [38]. Videos were only recorded during movement trials.

Muscle activity of the four compartments of the FDS (FDS2, FDS3, FDS4, FDS5) and EDC (EDC2, EDC3, EDC4, EDC5) were recorded using bipolar dual surface electrodes with a fixed interelectrode distance of 2 cm (SEMG/NCV Electrodes, Natus Neurology Inc., WI, USA). The electrode locations were based on published recommendations and experimental studies [1, 2, 39] and further guided by manual palpation. EMG signals were differentially amplified (CMRR > 115 dB, input impedance ∼ 10 GΩ) and bandpass filtered (10-1000 Hz) (AMT-8, Bortec Biomedical Ltd, AB, CA). EMG and force signals were sampled synchronously at 2000 Hz (16 bit, USB-6229, National Instruments, TX, USA) and collected using a custom-designed program (LabView 2016, National Instruments, TX, USA). The EMG recordings will not be addressed in this communication as any changes in muscle activity following fatigue could not be dissociated between alterations in muscle coordination versus myoelectric fatigue artefacts [40]. In addition, we observed substantial crosstalk between finger compartments upon visual inspection of the signals. Although some crosstalk is anticipated, the higher than expected in our study could be due to recruited sample differences between our study and previously conducted work, with individual differences in forearm anatomy and size known to affect crosstalk [39]. The EMG signals from the pre-fatigue isometric submaximal contractions are reported in the supplementary document (Figure S1).

### 2.3. Experimental Protocol

Participants visited the lab on two occasions each separated by at least 7 days. The experimental protocol across the visits were largely identical, involving a series of finger flexion-extension movements and isometric flexion or extension contractions at baseline and immediately following a fatigue protocol (Figure 2). The only difference across the two visits was whether the fatigue protocol targeted the ring flexors or ring extensors. The tasks performed by the participant during the baseline (i.e., pre-fatigue), fatigue protocol, and post-fatigue phases of the experimental protocol are described in the following sections.

**Figure 2:**
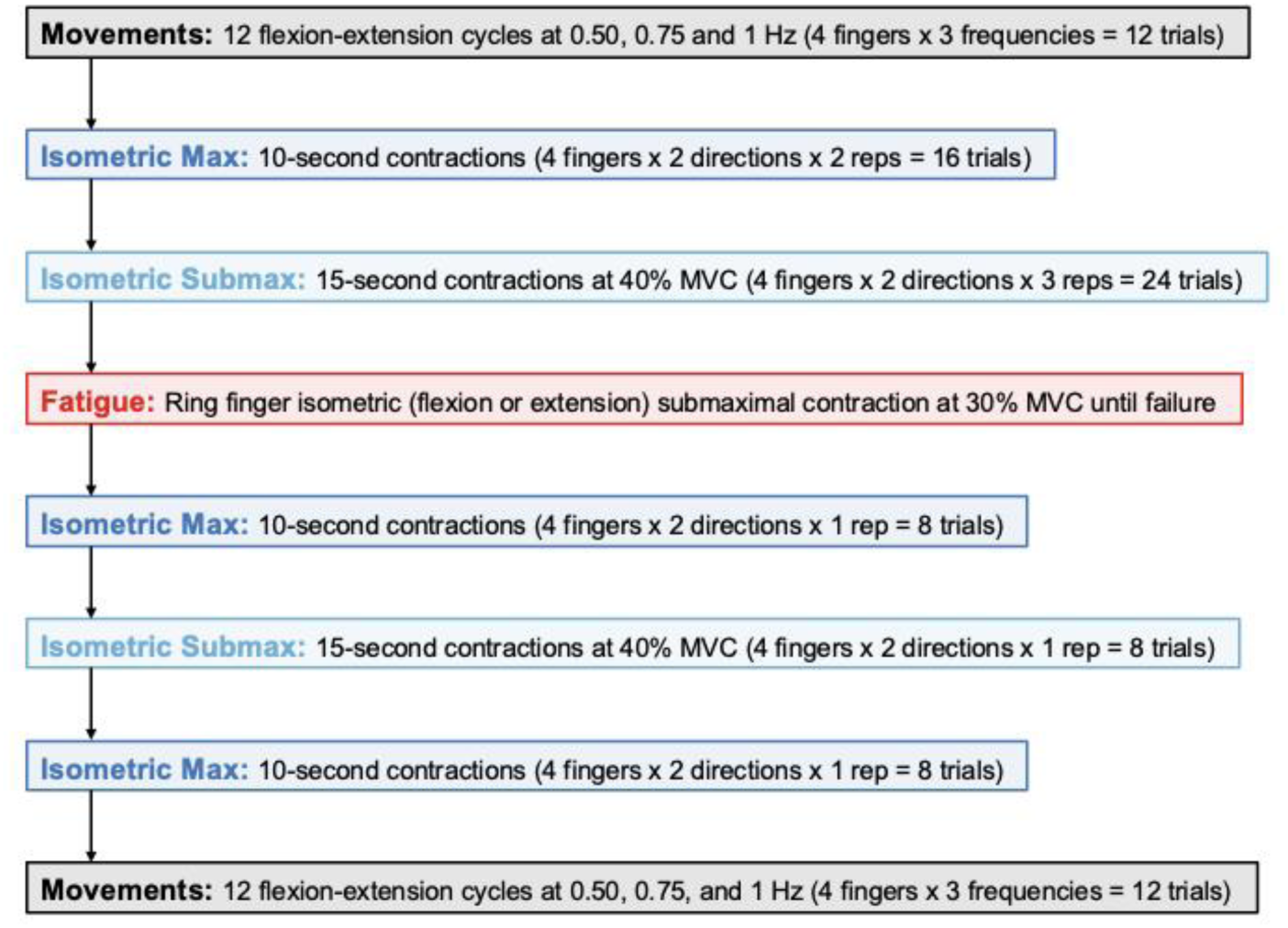
A flowchart of the experimental protocol. The protocol was repeated on two separate occasions, with the fatigue protocol (ring finger flexion or extension) randomized across the two visits. See main text for task descriptions.

#### 2.3.1. Baseline (Pre-Fatigue) Tasks

During finger movement trials, participants repetitively flexed and extended the instructed finger (index, middle, ring, or little) between two target postures. Starting from a straight (extended) finger, one flexion-extension cycle was defined as flexing the instructed finger to contact the palm (or as far as possible) and returning to a straight finger posture. Each trial involved 12 continuous flexion-extension cycles. The movements were completed at speeds of 0.50, 0.75, and 1 Hz (i.e., 60, 90, and 120 beats per minute) guided by auditory feedback from a metronome. A total of 12 movement trials (4 fingers x 3 speeds) were recorded. Trials were block randomized such that all movements of a single speed were completed in a randomized order before proceeding to the next speed (order of speeds was randomized). Participants were provided time to practice and familiarize themselves with the movements.

During isometric tasks, participants performed maximal and submaximal voluntary contractions with each instructed finger in either the flexion or extension direction. An initial 10 second quiet trial with the participant’s arm affixed in the apparatus was collected and used to debias the force and EMG signals. During the quiet trials, participants were instructed to relax and not actively exert with any of their fingers. For maximal voluntary contractions (MVC), participants were asked to exert force against the finger cuff ramping up to their maximum until instructed to relax. A total of 16 pre-fatigue MVC trials (4 fingers x 2 directions x 2 repetitions) were performed. The MVC trials were 10 seconds long and 1 minute of rest was provided between repetitions. A third MVC rep was performed if the peak force between the two repetitions for a given finger and direction combination was greater than 10%. The highest peak force across the repetitions was used as the MVC for each finger. For submaximal contractions, participants were instructed to maintain their finger force as closely as possible to a target level of 40% MVC (Figure 1b). Visual feedback of the instructed finger as well as a target force line was provided on a display monitor in front of the participants. A total of 24 pre-fatigue submaximal trials (4 fingers x 2 directions x 3 repetitions) were performed. The submaximal trials were 15 seconds long with 15 seconds of rest provided between trials. For both the isometric MVC and submaximal contractions (Figure 2), the order of trials were randomized across each finger and direction combination (i.e., all repetitions of a single combination were completed before proceeding to the next randomly determined combination). Participants were instructed to concentrate on producing force with only the instructed finger. In addition, participants were asked to maintain their trunk posture upright and avoid leaning into the apparatus to assist with force production.

#### 2.3.2. Fatigue Task

The fatiguing protocol involved an isometric submaximal contraction with the ring finger in either the flexion or extension direction. The exertion direction was randomized across the two visits. The target force level on both visits was set at 30% MVC. Participants continuously performed the isometric submaximal contraction until they reached one of the two termination criteria: (1) verbal declaration to discontinue, or (2) failure to maintain task performance as evident by the inability to reach the target force level despite verbal encouragement.

#### 2.3.3. Post-Fatigue Tasks

Immediately following the fatiguing protocol, participants repeated their baseline isometric contractions and movements. The order of the tasks was 8 isometric maximal contractions (4 fingers x 2 directions), 8 isometric submaximal contractions at 40% of baseline MVC (4 fingers x 2 directions), 8 isometric maximal contractions (4 fingers x 2 directions), and 12 movement trials (4 fingers x 3 frequencies). The set of maximal contractions were repeated twice to quantify strength declines immediately following the fatigue protocol (post-fatigue I) and possible strength recovery nearing the end of the experiment (post-fatigue II).

### 2.4. Data Processing and Analysis

Raw forces were de-biased and filtered (low-pass 2^nd^ order Butterworth, 4 Hz cut-off). All forces were normalized to baseline MVC. For isometric submaximal contractions, the root mean square difference between the measured and target force of the instructed finger was used to find the 5-second window where participants best matched the target force. The finger forces during the 5-second window were averaged, with the pre-fatigue finger forces additionally averaged across the three repetitions. In total, our isometric dataset comprised of finger forces from 960 isometric maximal trials (20 participants x 2 visits x 4 fingers x 2 directions x 3 fatigue phases [pre-fatigue, post-fatigue I, post-fatigue II]) and 640 isometric submaximal trials (20 participants x 2 visits x 4 instructed fingers x 2 directions x 2 fatigue phases [pre-fatigue, post-fatigue]). From these data, 5 maximal (0.5% of trials) and 10 submaximal (1.6% of trials) trials were removed due to participant exerting in the wrong direction with the instructed finger.

Kinematic data was analyzed using a markerless motion capture pipeline [38]. From the recorded videos, 21 key points across the hand were identified using MediaPipe [41] (Figure 1c). Extraneous points were removed using a median filter and interpolated with a cubic B-spline [42]. Camera synchronization was verified by performing a cross-correlation on the fingertip position of the instructed finger between camera pairs. The 2D positions from each camera were triangulated to obtain 3D positions [42] (Figure 1d). The 3D positions were filtered using a low-pass 2^nd^ order Butterworth filter (5 Hz cut-off). Joint angles of the MCP, PIP, and DIP were calculated as the angle between adjacent segments (e.g., MCP angle was the angle between the metacarpal [wrist to MCP key points] and proximal phalanx segments [MCP to PIP key points]). The sum of the MCP and PIP angles was used to represent finger motion [35] (Figure 1e). The DIP angle was not used due to occasional tracking inaccuracies of the fingertip position for the uninstructed fingers. Regardless, the MCP and PIP angles comprised most of the movement (> 90% of the total finger angular motion). The middle 9 flexion-extension cycles were identified for all fingers based on peaks and troughs in the instructed finger motion (e.g., index finger angles were used to identify movement cycles for all fingers during index flexion-extension tasks). The median of the 9 peaks was used to represent the overall finger motion (i.e., peak finger flexion) during the flexion-extension movements. In total, our dataset comprised of 960 movement trials (20 participants x 2 visits x 3 speeds x 2 fatigue phases x 4 instructed fingers) for each of the 4 fingers, resulting in 3840 finger motions. From these, 43 finger motions (1.2% of the data) were removed due to collection or analysis issues (e.g., participant did not return to a straight finger with each flexion-extension cycle, occlusion, and/or poor tracking). All data analyses were performed in Python (version 3.8).

### 2.5. Statistical Analysis

To determine whether the fatigue protocol was successful at targeting the ring finger, we evaluated differences in finger strength (isometric maximal contractions) with fatigue. A linear mixed-effects model was used to evaluate the main effect of fatigue (pre-fatigue vs. post-fatigue I vs. post-fatigue II) on finger strength. Separate models were generated for each experimental condition: instructed finger (index, middle, ring, little), visit (flexor fatigue, extensor fatigue), and direction (flexion, extension).

To verify that participants performed isometric submaximal contractions and flexion-extension movements consistently following the fatigue protocol relative to baseline, we compared the forces and motions by the *instructed* finger pre-vs. post-fatigue. This was accomplished using a paired t-test to evaluate the null hypothesis of no difference between pre-vs. post-fatigue and an equivalence test (two one-sided t-tests) to evaluate the null hypothesis that the difference in means is more extreme than an a priori effect size [43]. Based on within-subject variability during the pre-fatigue tasks, we set the a priori effect sizes to 1% MVC for the isometric submaximal tasks and 10° for movement tasks. For the isometric submaximal tasks, we performed the statistical tests on both the force magnitude and variability (using the standard deviation). For the finger movements, we performed the test on the peak finger flexion angle. Together, a non-significant paired t-test (do not reject null hypothesis that effect is equal to zero) and a significant equivalence test (reject null hypothesis that effect is greater than the a priori threshold) allows us to conclude that the pre- and post-fatigue means are “statistically equivalent”. In other words, the instructed finger performed the isometric submaximal and movement tasks within ±1% MVC (magnitude and variability) and ±10°, respectively, pre-vs. post-fatigue.

To determine changes in finger independence with fatigue, we evaluated differences in *involuntary* finger forces and motion (i.e., peak flexion) by the *non-instructed* fingers with fatigue. A linear mixed-effects model was used to evaluate the main effect of fatigue (pre-fatigue vs. post-fatigue) on *involuntary* forces and motion. Separate models were generated for each experimental condition: instructed finger (index, middle, ring, little), visit (flexor fatigue, extensor fatigue), and either direction (isometric tasks only: flexion, extension) or speed (movement tasks only: 0.50, 0.75, 1 Hz).

All model assumptions (linearity, normality and homogeneity of variance of residuals) were verified through visual inspection. For mixed-effects models, any statistically significant main effects of fatigue were followed-up by pairwise comparisons using Tukey’s adjusted p-values for multiple comparisons. An alpha value of 0.05 was set for all statistical tests. All statistical analyses were performed in R (version 4.3.2).

## 3. Results

### 3.1. Finger Strength (Maximal Exertions)

In this section, we describe statistically significant changes in finger strength following the fatigue protocol, with mean [95% confidence intervals (CI)] differences post-fatigue relative to pre-fatigue strength reported. Finger strengths across both visits are displayed in Figure 3. Detailed summary data and pairwise comparisons of finger strengths are reported in the supplementary document (Tables S1 and S2). For ease of communication, we will use the terms agonist and antagonist to refer to muscle groups with respect to the targeted fatigue direction. For example, the index, middle, ring, and little flexors will be considered as the agonists on the ring flexion fatigue visit and antagonists on the ring extension fatigue visit. However, we want to highlight that although muscles belonging to these groups may be agonists and antagonists with respect to the flexion-extension direction, the same cannot be said about their contributions to other degrees of freedom [44].

**Figure 3:**
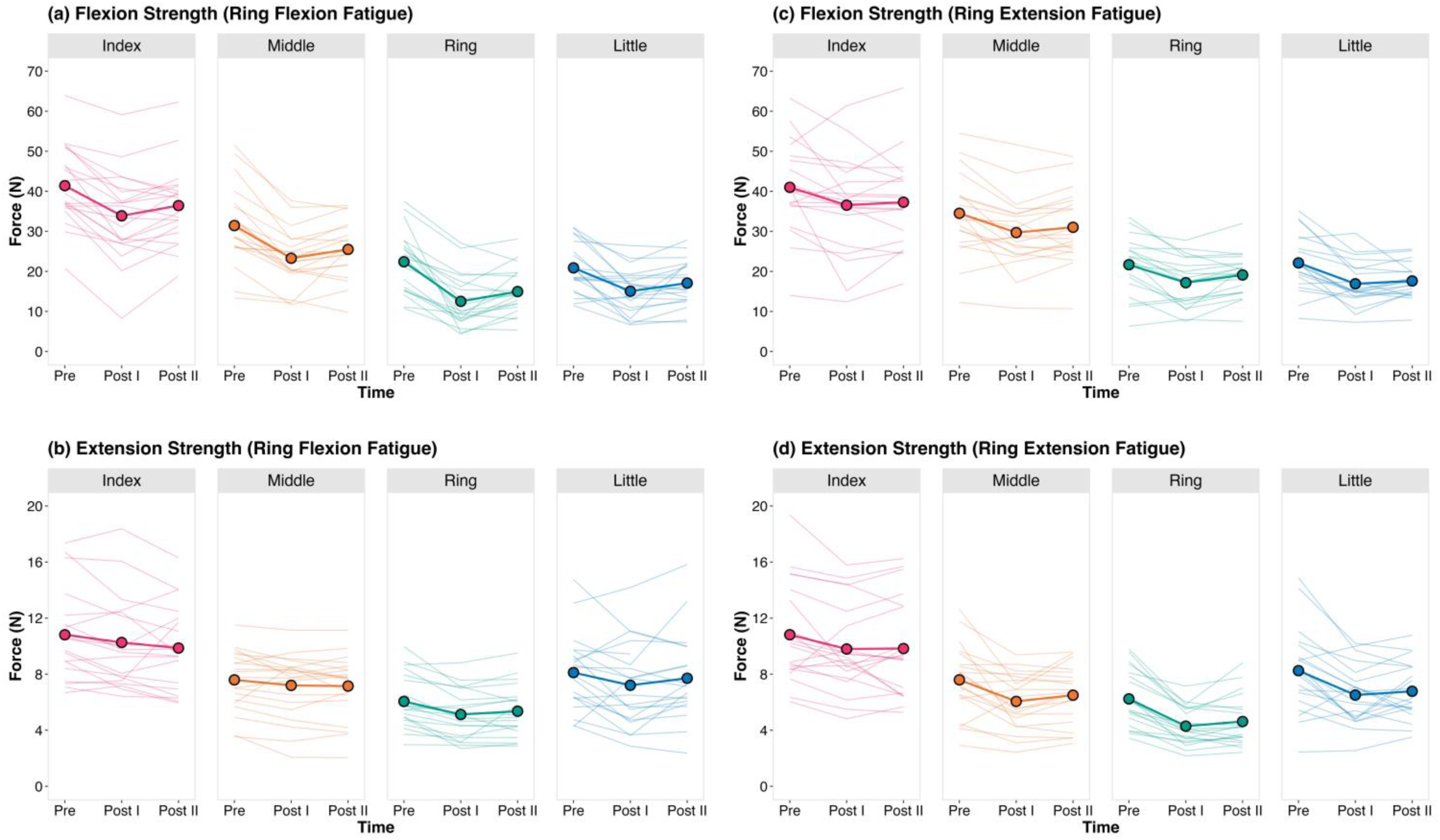
Changes in finger flexion and extension strength (N) with targeted fatigue of the ring finger flexors (a, b) and extensors (c, d). Strength measures were taken pre-fatigue (Pre), immediately following the fatigue protocol (Post I), and towards the end of the experiment prior to the post-fatigue movement tasks (Post II) (see Figure 2). The thick lines and points are group means, and the thinner lines are individual participant data. Please note the different y-axis scales for the finger flexion (a, c) and extension (b, d) strengths.

For the ring flexion and extension fatigue visits, we observed a decrease in ring flexion (−9.9 N [−12.8, −7.0]; Figure 3a) and extension strength (−1.9 N [−2.5, −1.4]; Figure 3d) respectively, immediately following the fatigue protocol. On average, the ring finger strengths were at 56.2% (flexion; range: 13.8-85.1%) and 70.0% (extension; range: 46.1-90.0%) relative to pre-fatigue values. Strength losses were concurrently observed across the ring finger antagonists for both visits. Ring extension strength decreased on the ring flexion fatigue visit (−0.9 N [−1.4, −0.5]; Figure 3b) and ring flexion strength decreased on the ring extension fatigue visit (−4.5 N [−6.5, - 2.5]; Figure 3c).

The non-ring fingers also experienced fatigue-induced strength deficits immediately following the fatigue protocol. For the ring finger flexion fatigue visit, flexion strength decreased across all other fingers (index: −7.5 N [−10.1, −4.9], middle: −8.2 N [−11.1, −5.3], little: - 5.8 N [−8.6, −3.1]; Figure 3a) but extension strength was not statistically different for any finger (index: −0.7 N [−1.6, 0.3], middle: −0.4 N [−0.9, 0.1], little: −0.9 N [−1.9, 0.1]; Figure 3b). For the ring extension fatigue visit, all non-ring fingers exhibited a reduction in flexion strength (index: - 4.5 N [−8.2, −0.7], middle: −4.8 N [−7.3, −2.3], little: −5.2 N [−7.5, −2.9]; Figure 3c) and extension strength (index: −1.0 N [−1.9, −0.1], middle: −1.5 N [−2.4, −0.7], little: −1.7 N [−2.7, −0.8]; Figure 3d).

All finger strength reductions immediately following the fatigue protocol (post-fatigue I) were also significantly lower during our second set of post-fatigue measures following the isometric submaximal tasks (post-fatigue II) relative to baseline. Although we did not find any statistically significant differences in finger strength between the two set of post-fatigue maximal contractions (post-fatigue 1 vs. post-fatigue II), strength values were not necessarily identical.

For instance, on the ring flexion fatigue visit, the 95% CI for the change in ring flexion strength between the post-fatigue measures (post-fatigue I vs. post-fatigue II) ranged from −0.4 N to 5.3 N. Thus, despite still exhibiting a strength deficit relative to baseline, a range of effects from further strength loss to some strength recovery were plausible throughout the post-fatigue phase of the experimental protocol.

### 3.2. Finger Independence: Isometric Submaximal Tasks

In this section, we first compare forces by the *instructed* finger pre-vs. post-fatigue to evaluate whether task performance was consistent with fatigue relative to baseline (also see Figure S2). We then compare *involuntary* forces by the *non-instructed* fingers pre-vs. post-fatigue to evaluate changes in finger independence with fatigue. Mean differences [95% CI] are expressed as post-fatigue relative to pre-fatigue (e.g., positive numbers indicate an increase in finger force with fatigue). Forces during isometric submaximal finger flexion (ring flexion fatigue visit) and extension (ring extension fatigue visit) tasks are displayed in Figures 4a-d and 5a-d, respectively. Forces during isometric tasks in the opposite direction of the targeted fatigue direction (i.e., finger extension tasks on the ring flexion fatigue visit, finger flexion tasks on the ring extension fatigue visit) are displayed in the supplementary document (Figure S3). Detailed summary data and pairwise comparisons of finger forces are also presented in the supplementary document (Tables S3-S6).

**Figure 4:**
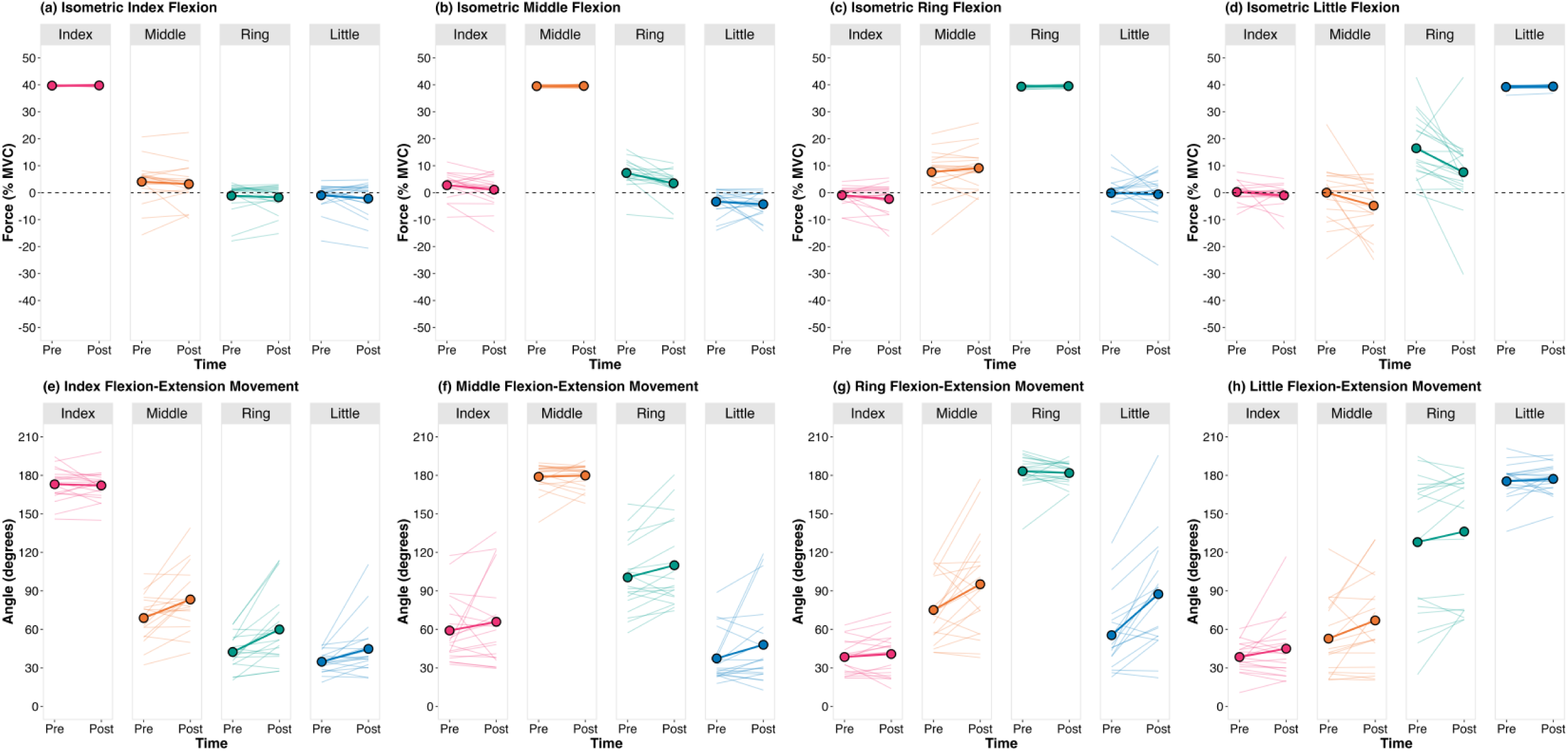
Changes in (a-d) finger forces (% MVC) during isometric submaximal (40% MVC) finger flexion and (e-h) peak flexion angles (°) during finger flexion-extension movements performed at 0.75 Hz with targeted fatigue of the ring finger flexors. The plot titles indicate the instructed finger. The thick lines and points are group means, and the thinner lines are individual participant data. Positive (and negative) values for the force data represent finger flexion (and extension) forces, respectively.

**Figure 5:**
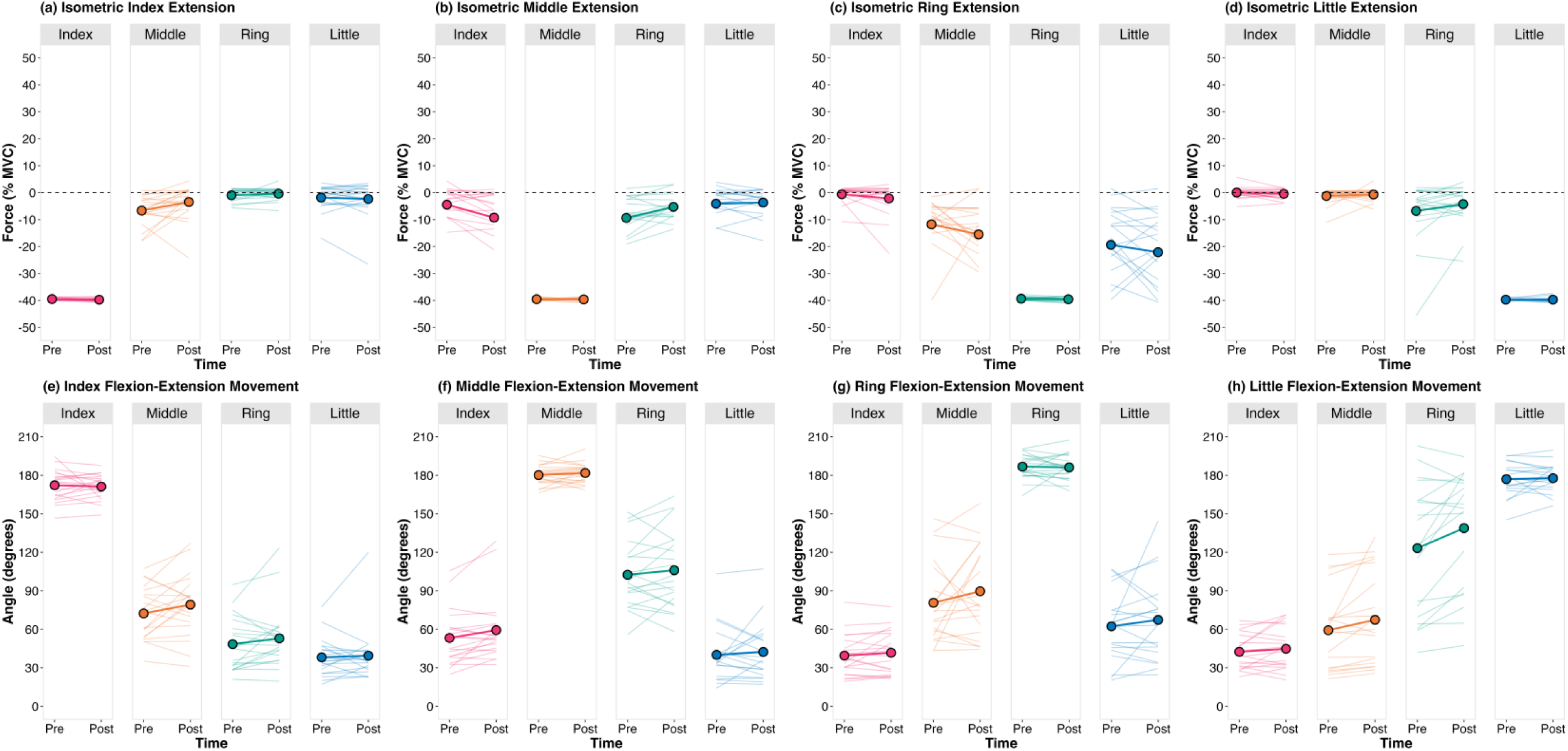
Changes in (a-d) finger forces (% MVC) during isometric submaximal (40%) finger extension and (e-h) peak flexion angles (°) during finger flexion-extension movements performed at 0.75 Hz with targeted fatigue of the ring finger extensors. The plot titles indicate the instructed finger. The thick lines and points are group means, and the thinner lines are individual participant data. Positive (and negative) forces represent finger flexion (and extension).

Using a paired t-test (null hypothesis significance test) and two one-sided tests (equivalence test), we confirmed that participants performed isometric submaximal force tasks consistently with their *instructed* finger pre-vs. post-fatigue (see instructed fingers in Figures 4a-d and 5a-d). Across most tasks, we found no significant difference in the magnitude of force produced by the *instructed* finger pre-vs. post-fatigue. The only exception was the index finger on the ring extension fatigue visit, where we observed a statistically significant increase in index finger force during index flexion (0.2 N [0.0, 0.4]; p=0.048; Figure S3a) and index extension (0.2 N [0.1, 0.4]; p=0.037; Figure 5a) post-fatigue relative to pre-fatigue. However, these differences are considered trivial as all equivalence tests were significant (p < 0.001), rejecting effects greater than our a priori bound of ±1% MVC. Thus, the true mean difference in the force produced by the *instructed* finger pre-vs. post-fatigue is within ±1% MVC. Similarly, significant equivalence tests confirmed that the difference in force variability of the *instructed* finger for all task conditions pre-vs. post-fatigue was within ±1% MVC. As a result, the magnitude and variability of the *instructed* finger force during the isometric submaximal tasks was equivalent pre-vs. post-fatigue.

Consistent with our a priori predictions, we found fatigue of the ring finger reduced *involuntary* ring finger forces during instructed exertions of adjacent fingers (middle and little). Following fatigue of the ring flexors, *involuntary* ring finger forces decreased during both middle finger flexion (−3.9 %MVC [−6.1, −1.7]; p=0.002; Figure 4b) and little finger flexion (−8.9 %MVC [−15.3, −2.5]; p=0.009; Figure 4d). Following fatigue of the ring extensors, involuntary ring finger forces decreased during middle finger extension (−2.6% MVC [−4.8, −0.4]; p=0.026; Figure 5b); however, we did not find statistically significant differences during little finger extension (−2.5% MVC [−5.7, 0.7]; p=0.113; Figure 5d). For these tasks (middle and little finger flexion and extension), the pre-fatigue involuntary ring finger forces ranged from 7.3-16.5% MVC when averaged across the group. Thus, the observed 95% CI (change in involuntary ring finger force with fatigue), which encompassed small positive effects (less than 1% MVC in magnitude) to large negative effects (greater than 5% MVC in magnitude) (see ring finger in Figures 4b, d and 5b, d), suggest our data is generally compatible with ring finger fatigue not affecting or decreasing *involuntary* ring finger forces. Outside our a priori predictions, we found no statistically significant differences in the involuntary finger forces across any of the other task conditions except for the following two instances: (i) increased involuntary ring finger force during middle finger extension following ring flexor fatigue (3.1% MVC [1.3, 4.8]; p=0.002; Figure S3f), and (ii) decreased involuntary middle finger force during index finger extension following ring extensor fatigue (−3.2% MVC [−6.2, −0.2]; p=0.040; Figure 5a).

### 3.3. Finger Independence: Flexion-Extension Movements

Finger motion during movement trials at the 0.75 Hz speed are displayed in Figures 4e-h and 5e-h for the ring flexion fatigue and extension fatigue visits, respectively. Figures for the remaining speeds (0.50 and 1 Hz) are displayed in the supplementary document (Figures S4 and S5) along with detailed summary data and pairwise comparisons of finger motion (Tables S7-S10).

Using a paired t-test (null hypothesis significance test) and two one-sided tests (equivalence test), we confirmed that participants performed flexion-extension movements consistently with their *instructed* finger pre-vs. post-fatigue (see instructed fingers in Figures 4e-h and 5e-h). Across most tasks, we found no significant difference in the peak finger flexion of the *instructed* finger pre-vs. post-fatigue. The only exception was the ring finger at the 1 Hz movement speed during the ring flexion fatigue visit, where we observed a statistically significant decrease in ring finger peak finger flexion post-fatigue (−5.5° [−7.9, −3.2]; p=0.001; Figure S4g). However, all equivalence tests were significant, allowing us to reject mean differences pre-vs. post-fatigue greater than our a priori bound of ±10°. Thus, the true mean differences in peak finger flexion of the *instructed* finger pre-vs. post-fatigue were within ±10°.

We predicted *a priori* that fatigue of the ring finger would not alter *involuntary* ring finger motion during instructed movements of adjacent fingers (middle and little). This prediction largely held for the ring flexion fatigue visit, where we didn’t find many statistically significant differences in *involuntary* ring motion, although large 95% CI in the positive direction (i.e., increased *involuntary* ring motion post-fatigue) were observed. During *instructed* middle finger movements, *involuntary* ring finger motion was significantly greater with fatigue for the 0.75 Hz task speed (9.4° [2.1, 16.6]; p=0.014; Figure 4f) but not significantly different at the 0.50 (6.6° [−0.6, 13.7]; p=0.069; Figure S4b) and 1 Hz (12.4° [−0.3, 25.2]; p=0.055; Figure S4f) speeds. During *instructed* little finger movements, *involuntary* ring finger motion was not significantly different following fatigue across all three task speeds: 0.50 (6.5° [−1.1, 14.0]; p=0.090; Figure S4d), 0.75 (6.3° [−1.9, 14.6]; p=0.123; Figure 4h), and 1 Hz (6.8° [−2.4, 15.9]; p=0.138; Figure S4h). On the ring extension fatigue visit, we tended to observe no changes or increased *involuntary* ring motion following fatigue. During *instructed* middle finger movements, *involuntary* ring finger motion was not statistically different at the 0.50 Hz (4.8° [−2.3, 12.0]; p=0.175; Figure S5b) and 0.75 Hz (3.5° [−4.3, 11.4]; p=0.357; Figure 5f) speeds but was significantly greater with fatigue at the 1 Hz speed (7.1° [2.8, 11.4]; p=0.003; Figure S5f).

During *instructed* little finger movements, *involuntary* ring finger motion was significantly greater with fatigue at the 0.50 Hz (9.3° [1.4, 17.1]; p=0.023; Figure S5d) and 0.75 Hz (14.4° [4.2, 24.5]; p=0.008; Figure 5h) speeds, but not significantly different for the 1 Hz task (5.3° [−3.2, 13.7]; p=0.206; Figure S5h). Outside our a priori predictions, we found either no statistically significant changes or significantly greater involuntary finger motion across the other task conditions. Across all our findings, the 95% CI encompassed small negative effects (less than 2 to 5° in magnitude) to large positive effects (greater than 10 to 20° in magnitude), suggesting our data is generally compatible with ring finger fatigue not affecting or increasing *involuntary* ring finger motion.

## 4. Discussion

This study sought to examine the relative contributions of neural and mechanical constraints of finger independence using a targeted fatigue protocol of the ring finger. Fatigue altered finger independence in opposing directions depending upon the task. Following fatigue, involuntary ring finger forces decreased while involuntary ring finger movement generally did not change or increased. Our results highlight that neuromechanical constraints to finger independence are fundamentally different for isometric versus movement tasks, which may be the source of confusion and conflicting literature on the roles of neural and mechanical factors in limiting finger independence.

The fatigue protocol successfully targeted the intended muscle group of the ring finger, but fatigue was not isolated. For both visits, we observed a decrease in strength of the ring finger agonists (i.e., ring finger flexion and extension strength decreased following targeted fatigue of the flexors and extensors, respectively). The mean strength declines across our participants ranged from approximately 30% (extension strength) to 45% (flexion strength) of pre-fatigue MVCs, which is in line with targeted fatigue of the index finger [45–47]. In addition, the agonists across all non-fatigued fingers exhibited strength declines (e.g., index, middle, little finger flexion strength decreased following targeted ring finger flexion fatigue). Although a formal statistical test was not conducted, the relative decrease in strength of the non-targeted fingers was less than the ring finger. Consistent with other studies [45–47], targeted fatigue of a single finger leads to strength deficits across agonists of all fingers. Based on our study, we can now generalize these findings to the ring finger and across contraction directions (flexion and extension). In addition, for the first time, we demonstrated that finger fatigue is also observed among antagonist muscles. For both visits, the ring finger antagonists displayed strength deficits (i.e., ring extension strength decreased during targeted fatigue of the ring flexors and vice versa). Further, for only the targeted ring extensor fatigue visit, we observed fatigue of the antagonist muscles across all non-targeted fingers (index, middle, little finger flexion strength decreased).

The latter result is especially surprising as antagonist muscle activity is typically higher during finger flexion tasks than extension [2, 37]. As the role of the finger and wrist extensors are primarily considered stabilizing and highly active during most hand tasks [48, 49], perhaps the extensors are less fatigable than the flexors, which aligns with the larger relative strength declines in the flexor (Figure 3a) compared to the extensor agonists (Figure 3d). Combined with existing evidence [45–47], the wide-spread strength deficit across all fingers highlights that isolated fatigue of a single finger during voluntary isometric contractions is not possible.

Targeted fatigue of the ring finger generally increased ring finger independence (i.e., decreased involuntary ring finger forces) during isometric tasks. Considering both the statistically significant differences and the range of plausible effects (95% CI), fatigue tended to decrease involuntary ring finger forces during instructed middle and little finger submaximal isometric tasks. While this result is consistent with our a priori predictions, which would support the hypothesis that neural factors are the primary constraints to finger independence during isometric contractions, the findings need to be interpreted in light of the complications arising from the widespread fatigue across fingers. Our predictions were made on the basis that a muscle will generate less force for the same amount of neural drive following fatigue [50]. If the middle and little fingers are not fatigued, we would expect the same amount of neural drive during instructed isometric contractions of these fingers pre-vs. post-fatigue of the ring finger.

This would result in the same diverging neural drive (pre-vs. post-fatigue) to the uninstructed ring finger and consequently, less involuntary ring finger forces with fatigue. Even if the middle and little fingers are fatigued, we assumed (and observed) that these fingers would not be as strongly fatigued as the ring finger based on earlier studies of the index finger [45–47]. In such a case, despite increased neural drive to the middle and little fingers to sustain the target force [51] leading to increased synaptic input to the uninstructed ring finger, the greater strength deficit of the ring finger would offset the increased neural drive and lead to a net decrease in involuntary ring forces. Our results are compatible with these explanations, with involuntary ring finger forces decreasing following targeted ring finger fatigue in the presence of the less fatigued middle and little fingers.

In contrast, targeted fatigue of the ring finger generally did not change or decreased ring finger independence (i.e., increased involuntary ring finger movement) during movement tasks. These results are in partial agreement with our a priori predictions (no change in involuntary ring finger movement). The increased involuntary ring finger movements were most prominent when following targeted fatigue of the ring finger extensors. As our movement outcome was peak finger flexion angle, the increase in involuntary ring finger movement post-fatigue can be explained by the weakened ring extensors. More curious are the findings from the ring finger flexion fatigue visit, where we saw plausible effects ranging from no change to increased involuntary ring finger movement. If neural factors played a substantial role in constraining finger independence during finger flexion movements, weakened ring flexors would decrease involuntary ring finger movement. Hence, our results could suggest that neural constraints to finger independence play a relatively minor role during finger movements. The tendency for involuntary ring finger movement to increase following fatigue of the ring flexors was unexpected. This is likely driven by the weakened ring extensors, but alternative mechanisms are plausible. One possibility is that descending pathways may contribute differently based on the motor task, which could shift with fatigue. The reticulospinal tract has greater divergence in synaptic input across muscles and is thought to have larger contributions during gross motor tasks such as grasping as opposed to the corticospinal tract, which is less divergent and contributes more towards fine motor skills requiring individual finger control [52–55]. Perhaps finger movements post-fatigue involves the reticulospinal tract to a greater extent, leading to increased divergent neural drive to the uninstructed ring finger and the recruitment of rested motor units not involved during isometric contractions, thereby increasing involuntary ring finger movement. It is also possible that increased involuntary movements may be due to impaired sensory feedback with fatigue [56]. As afferent signals can relay information not only about muscle length changes, but also relative changes in length between neighbouring muscles [57, 58], impaired feedback signals with fatigue could alter movement trajectories [59] of the uninstructed fingers. Consistent with the alterations in sensory feedback, several of our participants reported subjective feelings of stiffness and soreness during the post-fatigue movement trials, and it is possible that the increased involuntary movements helped minimize pain from relative strain in connective tissue linkages between fingers.

Together, our findings demonstrate that the effects of fatigue on finger independence were task-dependent, with opposite responses found for static versus movement tasks. These results add to the body of literature indicating that the neuromechanical control of force versus movement can be distinct with each other [60–62]. Yet, at least in the context of finger independence, treating neural versus mechanical factors as two mutually exclusive entities pitted against each other as we presented in the introduction (and as is mostly done in the literature) may be a naïve approach and can over-simplify the scientific problem. The neural and mechanical constraints can interact with one another in complex ways. As an example, divergent neural drive to a non-instructed finger will cause less relative displacement between musculotendon structures of adjacent fingers, thereby reducing strain of connective tissue linkages and decreasing passive transmission of forces. Furthermore, muscles do not act in isolation [44]. Muscles can act upon multiple kinematic degrees of freedom (e.g., extrinsic finger flexors exert moments around the wrist) that will require compensatory actions which can be a source of involuntary finger forces and movements [10]. The challenge in explaining the unexpected increase in involuntary finger movements we observed underscores the complexities of these neuromechanical constraints, which are speculated to have co-evolved over time [10]. Nevertheless, the opposing changes in finger independence across force and movement tasks exemplify that the contribution of neural and mechanical constraints depend on the task, and interpreting results across different studies must be made within the contexts of the experimental conditions. Moving forward, neuromusculoskeletal models of the fingers incorporating the plausible pathways for both neural and mechanical dependencies are required to tease apart the influence of these constraints under controlled, isolated simulations.

A key consideration in interpreting the results of our study are the limitations of the measurement outcomes. Uniaxial force transducers were used, allowing us to only measure finger flexion extension forces. Fatigue may have changed direction of finger force application, although this would have been inefficient as the target force was in the flexion-extension direction. In addition, our joint angle measurements do not separate out the components from different axes of rotation. Changes in involuntary movement may be due to small differences in abduction adduction angle rather than flexion-extension. Potential changes in finger force direction and abduction-adduction angle could be caused by altered muscle coordination patterns with increased use of intrinsic finger muscles post-fatigue. However, simulated muscle weakness at the wrist was previously found to simply scale up pre-existing muscle activity patterns rather than re-distribute loads across muscles [63]. This is also consistent with the highly structured solution space of muscle activity patterns during simulated finger contractions, with re-distributions in loads across finger muscles severely limited by biomechanical constraints [64]. Alternatively, any reduced contributions from neural constraints in a fatigued state would increase the relative change in length and position between adjacent musculotendon compartments. Thus, inter-connective tissue linkages will be stretched and pull the uninstructed fingers towards the instructed finger [12]. While we expect any changes in the direction of force application and abduction-adduction angle to be small as constraints to finger independence are primarily in the flexion-extension direction and due to task restrictions, future studies may warrant distinguishing between force and angle components.

In conclusion, targeted fatigue of the ring finger caused widespread fatigue across the entire hand, leading to generally decreased involuntary ring finger forces and no changes or increases in involuntary ring movement. By studying isometric finger tasks and movements under a single experimental paradigm, we demonstrated that finger independence can be altered in different ways based on task conditions. Thus, conflicting literature on the contributions of neural and mechanical constraints to finger independence may not necessarily be incompatible. Rather, differences in neuromechanical control across force and movement tasks can alter finger independencies. Our findings provide support to the hypothesis that involuntary forces during isometric tasks are limited primarily by neural constraints. In contrast, involuntary movements during finger movements are limited primarily by mechanical constraints. These differences naturally emerged from our nervous system responding to the differing task goals of isometric finger contractions versus movements under a fatigued condition.

### Ethics

The McMaster Research Ethics Board approved the study (MREB 6840).

### Use of Artificial Intelligence (AI) and AI-assisted technologies

We have not used any AI-assisted technologies in the preparation of our paper.

### Data, Code and Materials

All data and scripts required to replicate the study results are available open access (https://doi.org/10.17605/OSF.IO/EC29X) [65].

## CRediT Author Statement

- Daanish M. Mulla: Conceptualization, Methodology, Formal analysis, Investigation, Data Curation, Writing – Original Draft, Writing – Review & Editing, Visualization
- Paul M. Tilley: Methodology, Investigation, Data Curation, Writing – review & editing
- Peter J. Keir: Conceptualization, Methodology, Writing – review & editing, Supervision

## Acknowledgements

We would like to thank members of the McMaster Occupational Biomechanics Laboratory (Christian Cicco, Noelle Donatelli, and Joanna Misquitta) for their help in collecting the data.

