## Supplementary Information for "Unravelling neuromechanical constraints to finger independence"

### S1 Electromyography

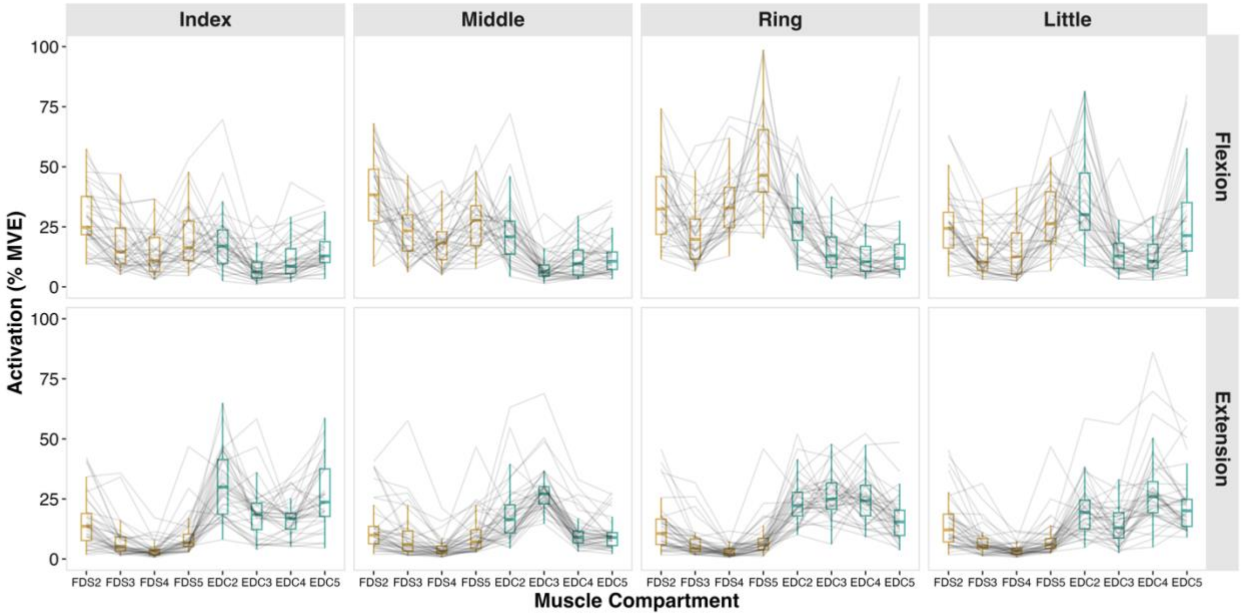


**Figure S1:** Mean EMG amplitude (% maximum voluntary exertion [MVE]) for all flexor digitorum superficialis (FDS2-FDS5) and extensor digitorum communis (EDC2-EDC5) compartments during pre-fatigue isometric submaximal tasks. The columns (index, middle, ring, little) denote the instructed finger. The rows (flexion, extension) denote the contraction direction. Coloured box and whisker plots represent the summary data (whiskers: minimum and maximum; boxes: 25th, 50th, and 75th percentile). The gray lines represent each participant. Data is compiled from both visits. The EMG amplitudes were calculated by debiasing, linear enveloping (low pass 2nd order Butterworth filter with a 4 Hz cut-off), and normalizing signals to the peak, filtered EMG amplitudes from the maximal contractions. The mean EMG amplitude was calculated as the average EMG amplitude during the 5-second window where participants best matched the target force (see Section 2.4).

**S2 Finger strengths**

**Table S1:** Summary data (mean ± standard deviation) of finger flexion and extension strength (N) across both visits prior to and following the fatigue protocol (pre-fatigue, post-fatigue I, and post-fatigue II).

| | **Ring Flexion Fatigue** | | | **Ring Extension Fatigue** | | |
|---|---|---|---|---|---|---|
| | Pre | Post I | Post II | Pre | Post I | Post II |
| **Flexion** | | | | | | |
| Index | 41.4±9.7 | 33.9±11.1 | 36.4±9.8 | 41.0±11.3 | 36.5±12.1 | 37.3±11.0 |
| Middle | 31.4±10.0 | 23.3±6.9 | 25.5±7.0 | 34.5±10.1 | 29.7±9.0 | 31.0±9.1 |
| Ring | 22.4±7.9 | 12.5±6.6 | 15.0±5.7 | 21.7±7.1 | 17.2±6.1 | 19.1±5.3 |
| Little | 20.9±6.3 | 15.0±5.6 | 17.1±5.7 | 22.1±7.1 | 16.9±5.6 | 17.6±4.4 |
| **Extension** | | | | | | |
| Index | 10.8±3.2 | 10.3±3.3 | 9.9±3.0 | 10.8±3.5 | 9.8±3.1 | 9.8±3.5 |
| Middle | 7.6±2.2 | 7.2±2.3 | 7.2±2.3 | 7.6±2.5 | 6.1±2.0 | 6.5±2.0 |
| Ring | 6.1±1.9 | 5.1±1.7 | 5.4±1.8 | 6.2±2.0 | 4.3±1.3 | 4.6±1.8 |
| Little | 8.1±2.7 | 7.2±3.0 | 7.7±3.2 | 8.3±3.0 | 6.5±2.1 | 6.8±2.0 |

**Table S2:** Linear mixed-effects model outputs evaluating the effects of fatigue on finger strength (N). The first column for each visit displays the F-statistics and p-values of the main effect of fatigue (pre-fatigue, post-fatigue I, post-fatigue II) on finger strength. The remaining columns display the means [95% confidence intervals] of the pairwise comparisons comparing across different levels of fatigue. For example, the value -7.5 N for index flexion under the column *Post I – Pre* on the ring flexion fatigue visit indicates that on average, there was a 7.5 N decrease in index flexion strength immediately following the fatigue protocol (Post I) compared to pre-fatigue (Pre) when targeting fatigue of the ring flexors.

| | **Ring Flexion Fatigue** | | | | **Ring Extension Fatigue** | | | |
|---|---|---|---|---|---|---|---|---|
| | F-stat, p | Post I – Pre | Post II – Pre | Post II – Post I | F-stat, p | Post I – Pre | Post II – Pre | Post II – Post I |
| **Flexion** | | | | | | | | |
| Index | 25.4 | -7.5 | -5.0 | 2.6 | 4.8 | -4.5 | -3.7 | 0.7 |
| | < 0.001 | [-10.1, -4.9] | [-7.6, -2.3] | [-0.1, 5.2] | 0.013 | [-8.2, -0.7] | [-7.5, 0.0] | [-3.0, 4.5] |
| Middle | 25.1 | -8.2 | -5.9 | 2.2 | 12.1 | -4.8 | -3.5 | 1.3 |
| | < 0.001 | [-11.1, -5.3] | [-8.8, -3.0] | [-0.7, 5.1] | < 0.001 | [-7.3, -2.3] | [-6.0, -1.0] | [-1.2, 3.8] |
| Ring | 37.7 | -9.9 | -7.4 | 2.4 | 15.0 | -4.5 | -2.6 | 1.9 |
| | < 0.001 | [-12.8, -7.0] | [-10.3, -4.6] | [-0.4, 5.3] | < 0.001 | [-6.5, -2.5] | [-4.6, -0.6] | [-0.1, 3.9] |
| Little | 14.2 | -5.8 | -3.8 | 2.0 | 18.5 | -5.2 | -4.5 | 0.7 |
| | < 0.001 | [-8.6, -3.1] | [-6.5, -1.1] | [-0.7, 4.8] | < 0.001 | [-7.5, -2.9] | [-6.8, -2.2] | [-1.6, 3.0] |
| **Extension** | | | | | | | | |
| Index | 3.2 | -0.7 | -0.9 | -0.3 | 5.0 | -1.0 | -1.0 | 0.0 |
| | 0.052 | [-1.6, 0.3] | [-1.9, 0.0] | [-1.2, 0.7] | 0.012 | [-1.9, -0.1] | [-1.9, -0.1] | [-0.9, 0.9] |
| Middle | 3.0 | -0.4 | -0.4 | 0.0 | 10.2 | -1.5 | -1.1 | 0.4 |
| | 0.064 | [-0.9, 0.1] | [-0.9, 0.0] | [-0.5, 0.4] | < 0.001 | [-2.4, -0.7] | [-2.0, -0.2] | [-0.4, 1.3] |
| Ring | 12.7 | -0.9 | -0.7 | 0.2 | 44.3 | -1.9 | -1.6 | 0.3 |
| | < 0.001 | [-1.4, -0.5] | [-1.2, -0.2] | [-0.2, 0.7] | < 0.001 | [-2.5, -1.4] | [-2.2, -1.1] | [-0.2, 0.9] |
| Little | 2.5 | -0.9 | -0.3 | 0.6 | 11.7 | -1.7 | -1.5 | 0.3 |
| | 0.098 | [-1.9, 0.1] | [-1.3, 0.7] | [-0.4, 1.6] | < 0.001 | [-2.7, -0.8] | [-2.4, -0.5] | [-0.7, 1.2] |

### S3 Submaximal Isometric Tasks

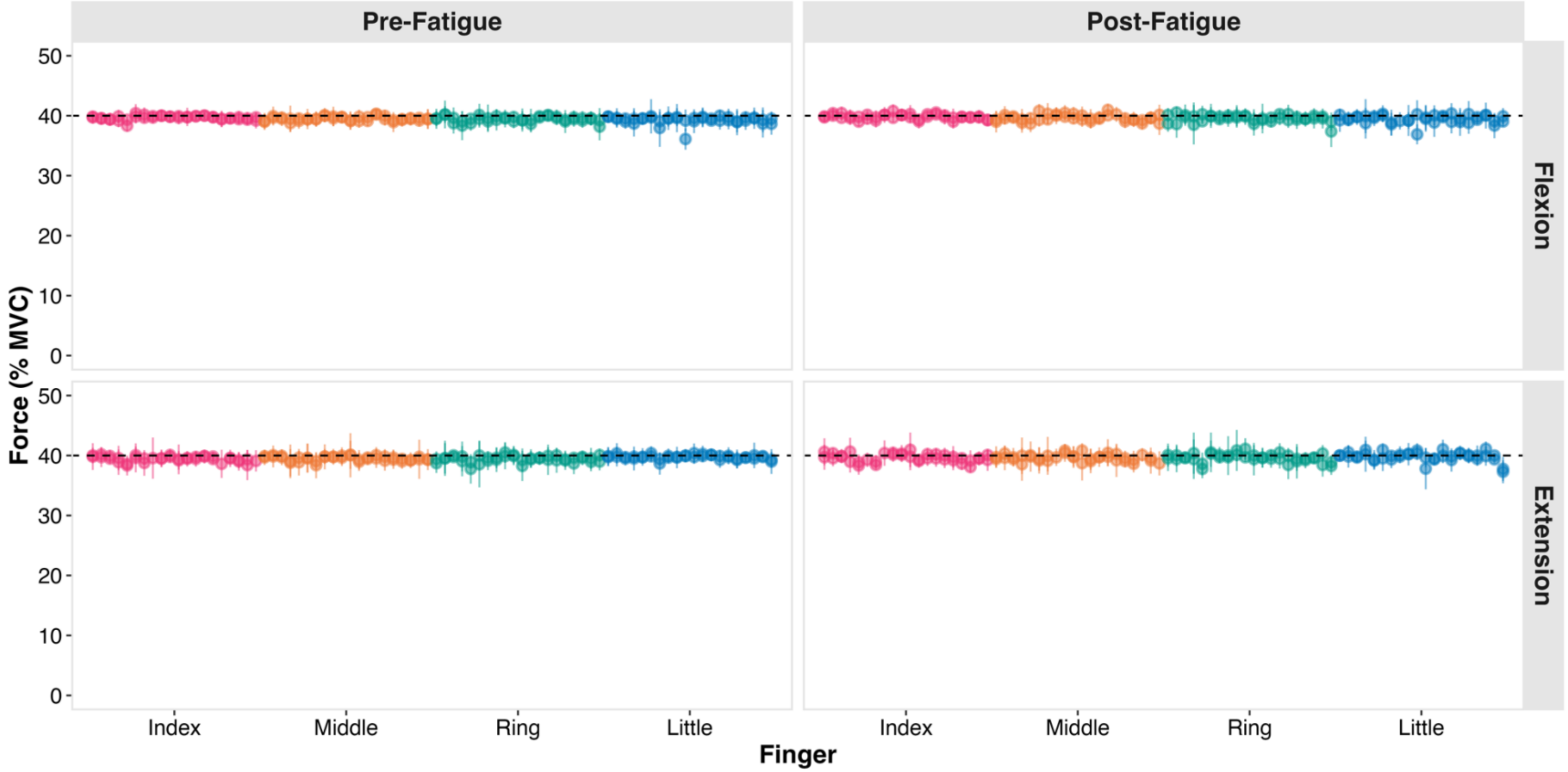


**Figure S2:** Forces (% MVC) by the *instructed* finger across all isometric submaximal tasks. The colours correspond to the instructed finger: index (pink), middle (orange), ring (green), and little (blue). Each point and error bar are a single participant's mean and standard deviation during the 5-second window where participants best matched the target force (40% MVC; horizontal dashed line). For pre-fatigue trials, data was pooled from the three repetitions. The points are ordered from left to right by participant, with data from both visits (ring flexion and extension fatigue) overlaid.

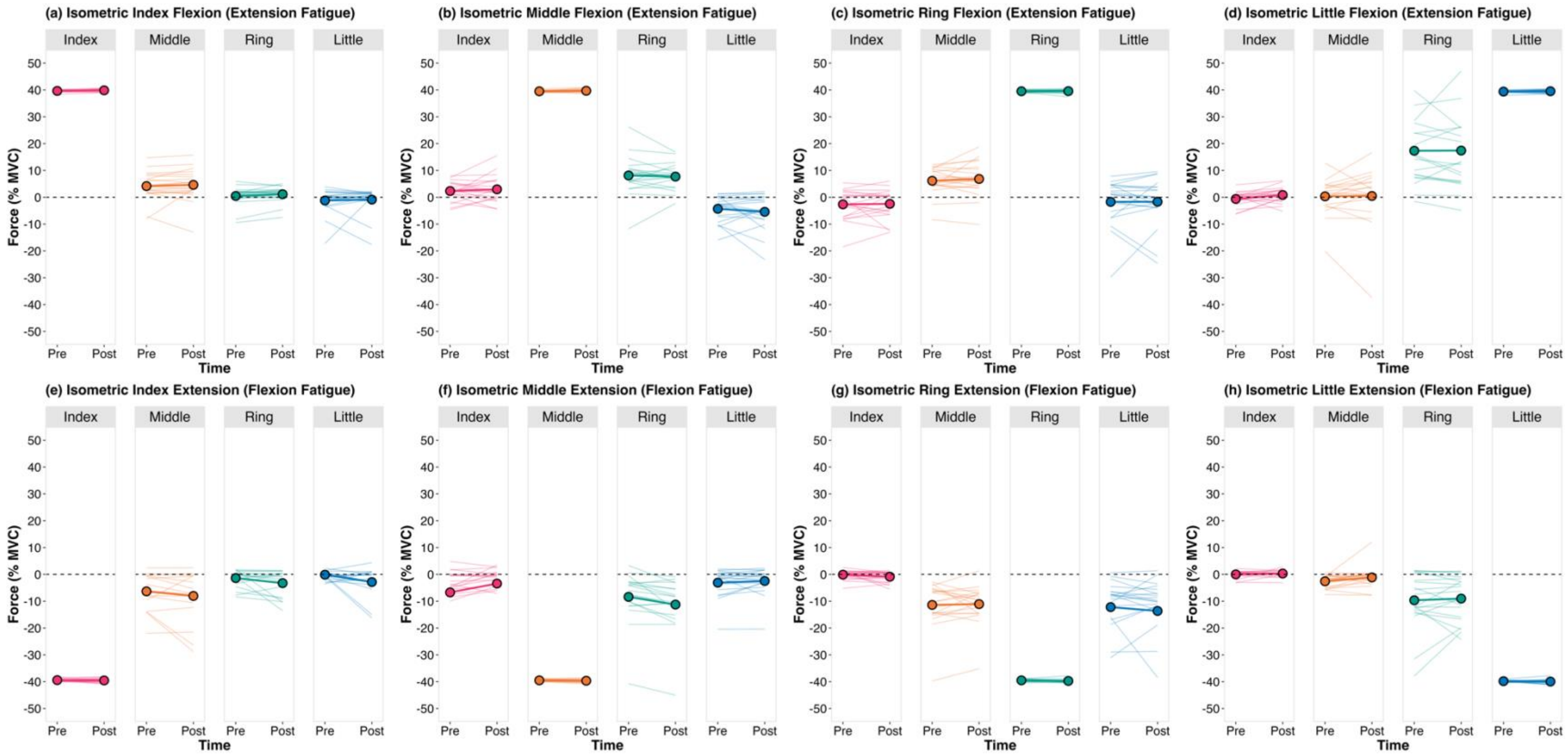


**Figure S3:** Finger forces (% MVC) by the muscle group antagonist to the muscles targeted during the fatigue protocol. (a-d) Changes in finger forces during isometric finger flexion following targeted fatigue of the ring finger extensors. (e-h) Changes in finger forces during isometric finger extension following targeted fatigue of the ring finger flexors. The plot titles indicate the instructed finger and exertion direction. The thick lines and points are group means and the thinner lines are individual participant data. Positive (and negative) forces represent finger flexion (and extension).

**Table S3:** Summary data (mean ± standard deviation) and pairwise comparisons pre- vs. post-fatigue (t-statistics, p-values, and estimated difference [95% confidence interval]) of finger forces (% MVC) across both visits during finger flexion tasks. For summary data, positive (negative) values indicate a finger force measured in the flexion (extension) direction. Estimates are based on linear mixed-effects model outputs and are expressed as post-fatigue relative to pre-fatigue. Please note that separate statistical tests were performed on the forces by the *instructed* fingers, hence no model outputs are presented in those cases (see Tables S5 and S6 instead).

| **Task** | **Ring Flexion Fatigue** | | | | | **Ring Extension Fatigue** | | | | |
|---|---|---|---|---|---|---|---|---|---|---|
| Measured Finger | Pre | Post | t | p | Estimate [95% CI] | Pre | Post | t | p | Estimate [95% CI] |
| **Index Flexion** | | | | | | | | | | |
| Index | 39.7±0.2 | 39.8±0.3 | | | | 39.6±0.4 | 39.9±0.5 | | | |
| Middle | 4.1±7.7 | 3.2±7.2 | -0.9 | 0.382 | -0.9 [-2.9, 1.2] | 4.2±5.4 | 4.6±6.0 | 0.6 | 0.529 | 0.5 [-1.0, 2.0] |
| Ring | -1.1±5.9 | -1.7±4.8 | -0.8 | 0.423 | -0.6 [-2.2, 1.0] | 0.5±3.6 | 1.1±2.9 | 2.1 | 0.054 | 0.7 [0.0, 1.3] |
| Little | -1.0±4.9 | -2.2±6.5 | -1.3 | 0.200 | -1.2 [-3.1, 0.7] | -1.2±4.7 | -0.9±4.9 | 0.3 | 0.772 | 0.3 [-2.1, 2.8] |
| **Middle Flexion** | | | | | | | | | | |
| Index | 2.8±4.6 | 1.1±5.7 | -1.7 | 0.099 | -1.6 [-3.6, 0.3] | 2.3±3.6 | 3.0±4.9 | 0.7 | 0.510 | 0.6 [-1.3, 2.6] |
| Middle | 39.5±0.4 | 39.6±0.5 | | | | 39.5±0.3 | 39.7±0.6 | | | |
| Ring | 7.3±4.9 | 3.5±5.1 | -3.7 | 0.002 | -3.9 [-6.1, -1.7] | 8.1±7.3 | 7.7±4.8 | -0.5 | 0.654 | -0.4 [-2.4, 1.5] |
| Little | -3.4±4.5 | -4.3±4.7 | -0.7 | 0.494 | -1.0 [-3.8, 1.9] | -4.3±4.9 | -5.4±6.5 | -1.0 | 0.353 | -1.1 [-3.6, 1.3] |
| **Ring Flexion** | | | | | | | | | | |
| Index | -0.9±3.5 | -2.3±5.7 | -1.4 | 0.190 | -1.4 [-3.6, 0.8] | -2.7±5.7 | -2.5±5.6 | 0.2 | 0.844 | 0.2 [-1.8, 2.2] |
| Middle | 7.6±8.0 | 9.1±7.1 | 1.4 | 0.178 | 1.5 [-0.7, 3.7] | 6.1±5.0 | 6.8±6.5 | 0.8 | 0.427 | 0.7 [-1.1, 2.4] |
| Ring | 39.3±0.5 | 39.6±0.5 | | | | 39.5±0.4 | 39.6±0.7 | | | |
| Little | -0.1±6.1 | -0.6±8.4 | -0.3 | 0.766 | -0.5 [-3.8, 2.8] | -1.7±8.8 | -1.7±9.1 | 0.0 | 0.979 | 0.0 [-3.2, 3.3] |
| **Little Flexion** | | | | | | | | | | |
| Index | 0.3±3.5 | -1.0±4.1 | -1.1 | 0.271 | -1.3 [-3.6, 1.1] | -0.6±2.8 | 0.8±2.9 | 2.0 | 0.057 | 1.4 [0.0, 2.9] |
| Middle | 0.0±10.1 | -4.8±9.3 | -2.1 | 0.051 | -4.8 [-9.7, 0.0] | 0.3±6.6 | 0.5±11.0 | 0.2 | 0.859 | 0.3 [-2.8, 3.3] |
| Ring | 16.5±11.4 | 7.6±13.4 | -2.9 | 0.009 | -8.9 [-15.3, -2.5] | 17.3±12.3 | 17.4±15.4 | 0.0 | 0.979 | 0.0 [-3.5, 3.5] |
| Little | 39.2±0.8 | 39.4±0.7 | | | | 39.4±0.4 | 39.5±0.6 | | | |

**Table S4:** Summary data (mean ± standard deviation) and pairwise comparisons pre- vs. post-fatigue (t-statistics, p-values, and estimated difference [95% confidence interval]) of finger forces (% MVC) across both visits during finger extension tasks. For summary data, positive (negative) values indicate a finger force measured in the flexion (extension) direction. Estimates are based on linear mixed-effects model outputs and are expressed as post-fatigue relative to pre-fatigue. Please note that separate statistical tests were performed on the forces by the *instructed* fingers, hence no model outputs are presented in those cases (see Tables S5 and S6 instead).

| **Task** | **Ring Flexion Fatigue** | | | | | **Ring Extension Fatigue** | | | | |
|---|---|---|---|---|---|---|---|---|---|---|
| Measured Finger | Pre | Post | t | p | Estimate [95% CI] | Pre | Post | t | p | Estimate [95% CI] |
| **Index Extension** | | | | | | | | | | |
| Index | -39.4±0.5 | -39.5±0.8 | | | | -39.5±0.4 | -39.7±0.6 | | | |
| Middle | -6.3±6.7 | -8.0±9.3 | -1.4 | 0.183 | -1.9 [-4.7, 1.0] | -6.7±5.5 | -3.5±6.2 | 2.2 | 0.040 | 3.2 [0.2, 6.2] |
| Ring | -1.4±3.6 | -3.3±4.9 | -1.7 | 0.115 | -1.9 [-4.3, 0.5] | -1.0±2.2 | -0.4±2.3 | 1.3 | 0.193 | 0.6 [-0.3, 1.5] |
| Little | -0.1±2.3 | -2.9±5.5 | -1.9 | 0.070 | -2.7 [-5.7, 0.2] | -1.8±4.5 | -2.3±6.5 | -0.7 | 0.500 | -0.5 [-2.1, 1.1] |
| **Middle Extension** | | | | | | | | | | |
| Index | -6.7±16.0 | -3.4±9.7 | 1.8 | 0.086 | 3.0 [-0.5, 6.5] | -4.5±7.9 | -3.9±14.8 | -1.4 | 0.171 | -5.0 [-12.4, 2.3] |
| Middle | -39.5±0.4 | -39.7±0.6 | | | | -39.5±0.4 | -39.6±0.6 | | | |
| Ring | -8.4±9.3 | -11.3±9.8 | -3.7 | 0.002 | -3.1 [-4.8, -1.3] | -9.4±10.8 | -5.3±4.6 | 2.5 | 0.026 | 2.6 [0.4, 4.8] |
| Little | -3.1±5.2 | -2.5±5.1 | 0.9 | 0.373 | 0.6 [-0.8, 2.0] | -4.1±6.6 | -3.7±4.9 | -0.8 | 0.428 | -0.6 [-2.3, 1.0] |
| **Ring Extension** | | | | | | | | | | |
| Index | -0.1±1.7 | -0.9±2.0 | -1.7 | 0.098 | -0.8 [-1.7, 0.2] | -0.6±2.9 | -2.1±6.0 | -1.4 | 0.183 | -1.6 [-3.9, 0.8] |
| Middle | -11.4±8.2 | -11.1±7.2 | 0.3 | 0.769 | 0.3 [-1.9, 2.5] | -11.8±10.9 | -15.5±13.6 | -1.5 | 0.149 | -3.7 [-8.9, 1.5] |
| Ring | -39.5±0.5 | -39.7±0.6 | | | | -39.3±0.6 | -39.5±0.8 | | | |
| Little | -12.2±11.0 | -13.6±15.2 | -0.7 | 0.464 | -1.4 [-5.5, 2.6] | -19.4±15.3 | -22.1±16.2 | -1.2 | 0.264 | -2.7 [-7.7, 2.2] |
| **Little Extension** | | | | | | | | | | |
| Index | 0.0±1.3 | 0.3±1.2 | 1.0 | 0.331 | 0.3 [-0.3, 1.0] | 0.0±2.1 | -0.4±1.7 | -1.2 | 0.236 | -0.5 [-1.3, 0.3] |
| Middle | -2.6±2.3 | -1.1±3.9 | 1.9 | 0.069 | 1.4 [-0.1, 3.0] | -1.2±2.8 | -0.7±2.2 | 0.7 | 0.480 | 0.6 [-1.1, 2.2] |
| Ring | -9.6±10.5 | -9.0±8.3 | 0.4 | 0.677 | 0.6 [-2.5, 3.7] | -6.8±11.1 | -4.3±7.3 | 1.7 | 0.113 | 2.5 [-0.7, 5.7] |
| Little | -39.8±0.4 | -40.0±0.8 | | | | -39.7±0.4 | -39.7±0.8 | | | |

**Table S5:** Statistical results (t-statistics, degrees of freedom, p-values, raw estimates [90% confidence interval], and Hedge's g effect size [90% confidence interval]) from the null hypothesis significance tests and equivalence tests comparing the force magnitude (% MVC) by the *instructed* finger pre- vs. post-fatigue. The effect sizes are expressed as post-fatigue relative to pre-fatigue (i.e., positive values indicate greater force post-fatigue vs. pre-fatigue). The a priori effect size for the equivalence tests was set at 1% MVC. Together, a non-significant null hypothesis test and a significant equivalence test allows us to conclude that the pre- and post-fatigue force magnitudes are equivalent (mean difference within ±1% MVC).

| | **Null hypothesis significance test** | | | **Equivalence test** | | | **Effect sizes** | |
|---|---|---|---|---|---|---|---|---|
| | t | df | p | t | df | p | Raw Estimate [90% CI] | Hedge's g [90% CI] |
| ***Ring Flexion Fatigue*** | | | | | | | | |
| **Flexion** | | | | | | | | |
| Index | 0.8 | 19 | 0.456 | -10.2 | 19 | < 0.001 | 0.1 [-0.1, 0.2] | 0.2 [-0.2, 0.5] |
| Middle | 1.2 | 18 | 0.257 | -10.8 | 18 | < 0.001 | 0.1 [0.0, 0.2] | 0.3 [-0.1, 0.6] |
| Ring | 1.5 | 19 | 0.160 | -5.8 | 19 | < 0.001 | 0.2 [0.0, 0.4] | 0.3 [-0.1, 0.7] |
| Little | 1.4 | 19 | 0.180 | -6.6 | 19 | < 0.001 | 0.2 [0.0, 0.4] | 0.3 [-0.1, 0.7] |
| **Extension** | | | | | | | | |
| Index | 0.6 | 15 | 0.532 | -4.9 | 15 | < 0.001 | 0.1 [-0.2, 0.4] | 0.2 [-0.2, 0.5] |
| Middle | 1.5 | 18 | 0.152 | -9.7 | 18 | < 0.001 | 0.1 [0.0, 0.3] | 0.3 [0.0, 0.7] |
| Ring | 1.6 | 19 | 0.136 | -5.5 | 19 | < 0.001 | 0.2 [0.0, 0.5] | 0.3 [0.0, 0.7] |
| Little | 0.9 | 19 | 0.401 | -5.0 | 19 | < 0.001 | 0.1 [-0.1, 0.4] | 0.2 [-0.2, 0.5] |
| ***Ring Extension Fatigue*** | | | | | | | | |
| **Flexion** | | | | | | | | |
| Index | 2.1 | 19 | 0.048 | -7.4 | 19 | < 0.001 | 0.2 [0.0, 0.4] | 0.5 [0.1, 0.8] |
| Middle | 1.6 | 19 | 0.125 | -6.9 | 19 | < 0.001 | 0.2 [0.0, 0.4] | 0.3 [0.0, 0.7] |
| Ring | 0.3 | 19 | 0.756 | -6.5 | 19 | < 0.001 | 0.0 [-0.2, 0.3] | 0.1 [-0.3, 0.4] |
| Little | 1.1 | 18 | 0.277 | -7.2 | 18 | < 0.001 | 0.1 [-0.1, 0.3] | 0.2 [-0.1, 0.6] |
| **Extension** | | | | | | | | |
| Index | 2.2 | 19 | 0.037 | -7.3 | 19 | < 0.001 | 0.2 [0.1, 0.4] | 0.5 [0.1, 0.9] |
| Middle | 0.4 | 17 | 0.663 | -5.1 | 17 | < 0.001 | 0.1 [-0.2, 0.4] | 0.1 [-0.3, 0.5] |
| Ring | 1.2 | 19 | 0.232 | -4.9 | 19 | < 0.001 | 0.2 [-0.1, 0.5] | 0.3 [-0.1, 0.6] |
| Little | 0.0 | 19 | 0.987 | 5.3 | 19 | < 0.001 | 0.0 [-0.3, 0.3] | 0.0 [-0.4, 0.3] |

**Table S6:** Statistical results (t-statistics, degrees of freedom, p-values, raw estimates [90% confidence interval], and Hedge's g effect size [90% confidence interval]) from the null hypothesis significance tests and equivalence tests comparing the force variability (% MVC) of the *instructed* finger pre- vs. post-fatigue. The effect sizes are expressed as post-fatigue relative to pre-fatigue (i.e., positive values indicate greater force variability post-fatigue vs. pre-fatigue). The a priori effect size for the equivalence tests was set at 1% MVC. Together, a non-significant null hypothesis test and a significant equivalence test allows us to conclude that the pre- and post-fatigue force variability are equivalent (mean difference within ±1% MVC).

| | **Null hypothesis significance test** | | | **Equivalence test** | | | **Effect sizes** | |
|---|---|---|---|---|---|---|---|---|
| | t | df | p | t | df | p | Raw Estimate [90% CI] | Hedge's g [90% CI] |
| ***Ring Flexion Fatigue*** | | | | | | | | |
| **Flexion** | | | | | | | | |
| Index | -0.9 | 19 | 0.394 | 18.3 | 19 | < 0.001 | 0.0 [-0.1, 0.0] | -0.2 [-0.5, 0.2] |
| Middle | 0.3 | 18 | 0.767 | -11.6 | 18 | < 0.001 | 0.0 [-0.1, 0.2] | 0.1 [-0.3, 0.4] |
| Ring | 1.5 | 19 | 0.149 | -7.2 | 19 | < 0.001 | 0.2 [0.0, 0.4] | 0.3 [0.0, 0.7] |
| Little | 1.0 | 19 | 0.329 | -7.7 | 19 | < 0.001 | 0.1 [-0.1, 0.3] | 0.2 [-0.1, 0.6] |
| **Extension** | | | | | | | | |
| Index | 0.0 | 15 | 0.974 | 6.3 | 15 | < 0.001 | 0.0 [-0.3, 0.3] | 0.0 [-0.4, 0.4] |
| Middle | -0.4 | 18 | 0.721 | 5.4 | 18 | < 0.001 | -0.1 [-0.4, 0.2] | -0.1 [-0.4, 0.3] |
| Ring | -1.1 | 19 | 0.289 | 7.3 | 19 | < 0.001 | -0.1 [-0.3, 0.1] | -0.2 [-0.6, 0.1] |
| Little | 0.4 | 19 | 0.730 | -9.4 | 19 | < 0.001 | 0.0 [-0.1, 0.2] | 0.1 [-0.3, 0.4] |
| ***Ring Extension Fatigue*** | | | | | | | | |
| **Flexion** | | | | | | | | |
| Index | -0.9 | 19 | 0.366 | 19.1 | 19 | < 0.001 | 0.0 [-0.1, 0.0] | -0.2 [-0.6, 0.2] |
| Middle | -1.0 | 19 | 0.326 | 14.9 | 19 | < 0.001 | -0.1 [-0.2, 0.0] | -0.2 [-0.6, 0.1] |
| Ring | -1.4 | 19 | 0.170 | 8.6 | 19 | < 0.001 | -0.1 [-0.3, 0.0] | -0.3 [-0.7, 0.1] |
| Little | -0.1 | 18 | 0.943 | 6.5 | 18 | < 0.001 | 0.0 [-0.3, 0.3] | 0.0 [-0.4, 0.3] |
| **Extension** | | | | | | | | |
| Index | -1.5 | 19 | 0.163 | 7.4 | 19 | < 0.001 | -0.2 [-0.4, 0.0] | -0.3 [-0.7, 0.1] |
| Middle | 4.0 | 17 | 0.001 | -5.8 | 17 | < 0.001 | 0.4 [0.2, 0.6] | 0.9 [0.4, 1.3] |
| Ring | 1.7 | 19 | 0.098 | -5.0 | 19 | < 0.001 | 0.3 [0.0, 0.5] | 0.4 [0.0, 0.7] |
| Little | 1.3 | 19 | 0.200 | -5.1 | 19 | < 0.001 | 0.2 [-0.1, 0.5] | 0.3 [-0.1, 0.6] |

### S4 Movement Tasks

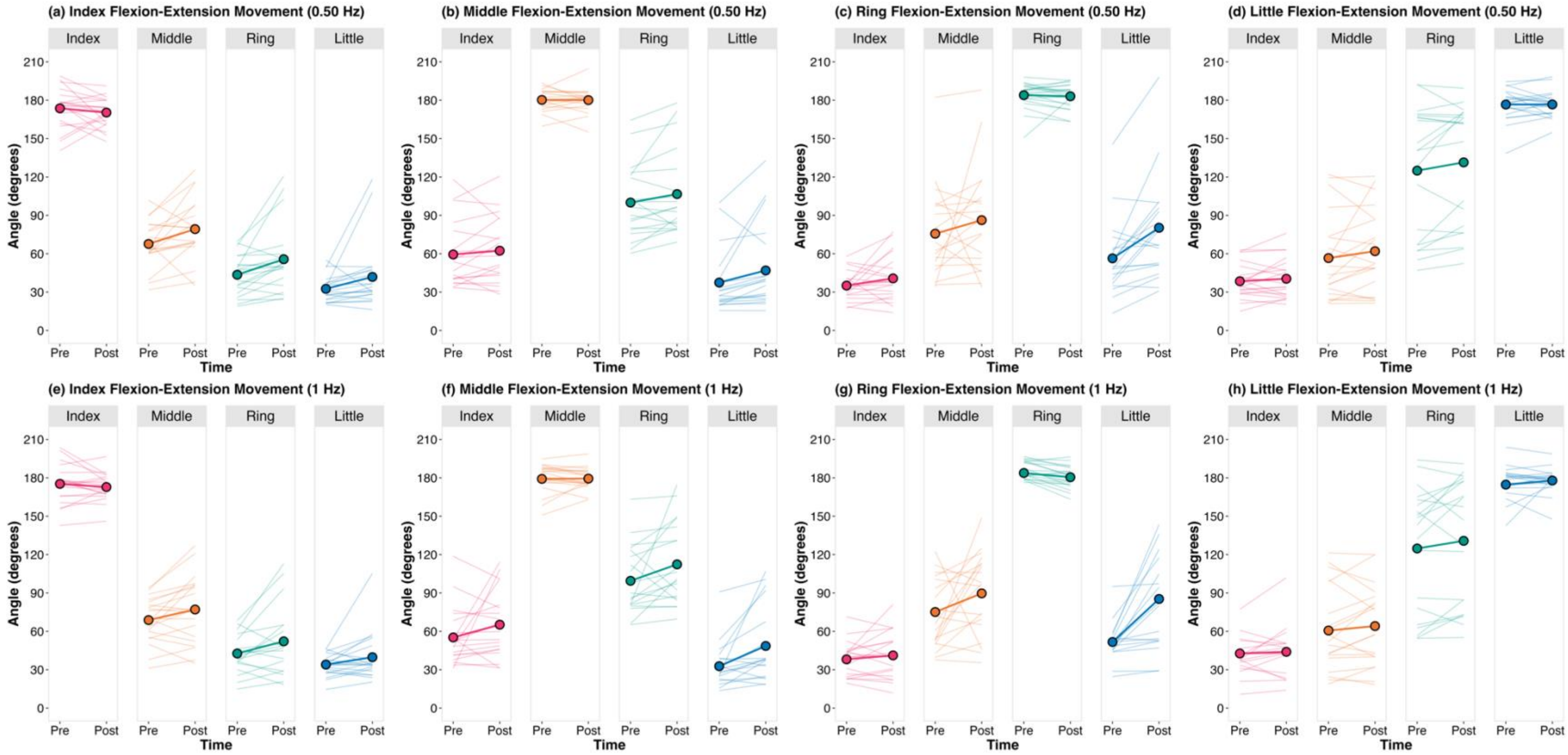


**Figure S4:** Changes in peak finger flexion angles (°) during flexion-extension movements performed at (a-d) 0.50 Hz and (e-h) 1 Hz with targeted fatigue of the ring finger flexors. The plot titles indicate the instructed finger and exertion direction. The thick lines and points are group means and the thinner lines are individual participant data.

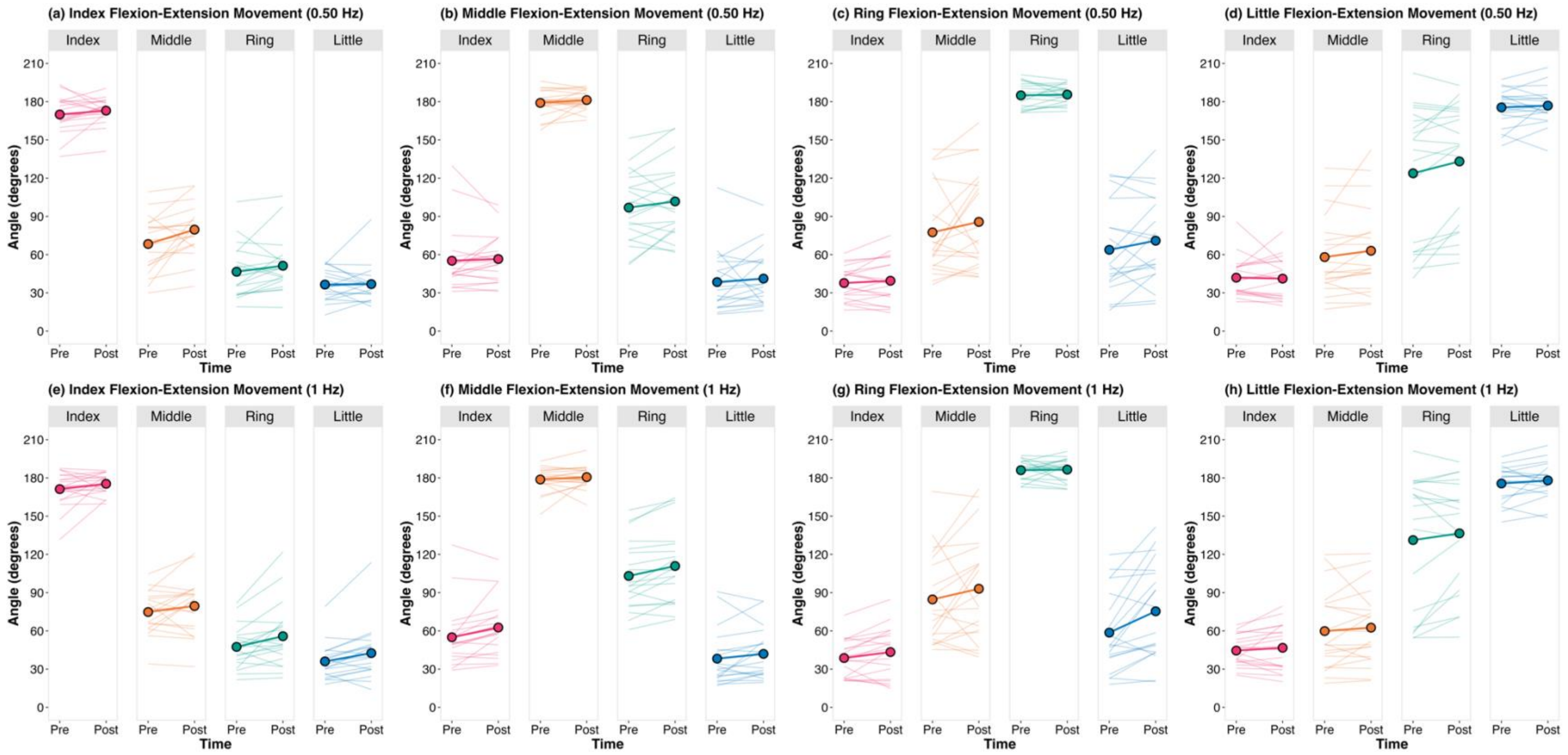


**Figure S5:** Changes in peak finger flexion angles (°) during flexion-extension movements performed at (a-d) 0.50 Hz and (e-h) 1 Hz with targeted fatigue of the ring finger extensors. The plot titles indicate the instructed finger and exertion direction. The thick lines and points are group means and the thinner lines are individual participant data.

**Table S7** Summary data (mean ± standard deviation) and pairwise comparisons pre- vs. post-fatigue (t-statistics, p-values, and estimated difference [95% confidence interval]) of peak flexion angle (°) across both visits during finger flexion-extension movements at 0.50 Hz.  Estimates are based on linear mixed-effects model outputs and are expressed as post-fatigue relative to pre-fatigue.  Please note that separate statistical tests were performed on the angles of the *instructed* fingers, hence no model outputs are presented in those cases (see Table S10 instead).

| **Instructed Finger** | **Ring Flexion Fatigue** | | | | | **Ring Extension Fatigue** | | | | |
|---|---|---|---|---|---|---|---|---|---|---|
| Measured Finger | Pre | Post | t | p | Estimate [95% CI] | Pre | Post | t | p | Estimate [95% CI] |
| **Index** | | | | | | | | | | |
| Index | 173.7±15.8 | 170.3±11.5 | | | | 169.9±14.3 | 173.0±10.2 | | | |
| Middle | 67.6±18.4 | 79.2±24.2 | 2.5 | 0.024 | 11.4 [1.7, 21.1] | 68.3±22.3 | 79.6±19.6 | 2.9 | 0.010 | 11.0 [2.9, 19.1] |
| Ring | 43.6±15.8 | 55.7±27.0 | 2.9 | 0.010 | 12.1 [3.2, 21.0] | 46.6±20.6 | 51.3±20.6 | 1.6 | 0.123 | 4.7 [-1.4, 10.7] |
| Little | 32.5±10.1 | 41.9±26.2 | 1.8 | 0.083 | 9.3 [-1.3, 20.0] | 36.5±11.9 | 36.9±14.5 | 0.1 | 0.894 | 0.4 [-5.5, 6.2] |
| **Middle** | | | | | | | | | | |
| Index | 59.4±25.8 | 62.4±25.1 | 1.0 | 0.351 | 3.0 [-3.6, 9.6] | 55.2±24.8 | 56.5±18.6 | 0.5 | 0.651 | 1.3 [-4.8, 7.5] |
| Middle | 180.2±8.4 | 180.1±9.7 | -0.2 | | | 179.0±10.3 | 181.3±7.5 | | | |
| Ring | 100.0±28.9 | 106.6±32.6 | 1.9 | 0.069 | 6.6 [-0.6, 13.7] | 96.9±27.2 | 101.7±29.1 | 1.4 | 0.175 | 4.8 [-2.3, 12.0] |
| Little | 37.5±24.7 | 46.9±32.8 | 2.2 | 0.039 | 10.4 [0.6, 20.2] | 38.5±23.6 | 41.2±21.9 | 0.8 | 0.454 | 2.7 [-4.7, 10.2] |
| **Ring** | | | | | | | | | | |
| Index | 35.1±11.2 | 40.7±17.9 | 1.6 | 0.123 | 5.6 [-1.7, 12.9] | 37.7±13.1 | 39.5±17.5 | 0.9 | 0.357 | 1.8 [-2.2, 5.7] |
| Middle | 75.6±35.7 | 86.2±41.0 | 1.1 | 0.295 | 10.0 [-9.5, 29.5] | 77.5±34.8 | 85.7±38.3 | 1.3 | 0.206 | 8.2 [-4.9, 21.3] |
| Ring | 183.9±10.6 | 183.1±9.7 | | | | 184.9±9.6 | 185.6±7.1 | | | |
| Little | 56.4±29.7 | 80.3±40.8 | 3.8 | 0.001 | 21.8 [9.8, 33.9] | 63.8±35.0 | 70.9±34.4 | 1.6 | 0.124 | 7.1 [-2.1, 16.4] |
| **Little** | | | | | | | | | | |
| Index | 38.5±13.7 | 40.4±15.6 | 1.0 | 0.336 | 1.9 [-2.2, 6.0] | 41.9±15.1 | 41.2±16.0 | -0.3 | 0.782 | -0.8 [-6.4, 4.9] |
| Middle | 56.6±33.2 | 62.0±32.5 | 1.3 | 0.207 | 5.4 [-3.2, 14.0] | 58.1±31.0 | 63.0±34.4 | 1.7 | 0.113 | 4.9 [-1.3, 11.2] |
| Ring | 125.0±50.7 | 131.4±47.7 | 1.8 | 0.090 | 6.5 [-1.1, 14.0] | 123.8±53.6 | 133.1±47.4 | 2.5 | 0.023 | 9.3 [1.4, 17.1] |
| Little | 176.6±12.8 | 176.7±10.2 | | | | 175.5±14.3 | 176.9±14.2 | | | |

**Table S8** Summary data (mean ± standard deviation) and pairwise comparisons pre- vs. post-fatigue (t-statistics, p-values, and estimated difference [95% confidence interval]) of peak flexion angle (°) across both visits during finger flexion-extension movements at 0.75 Hz. Estimates are based on linear mixed-effects model outputs and are expressed as post-fatigue relative to pre-fatigue. Please note that separate statistical tests were performed on the angles of the *instructed* fingers, hence no model outputs are presented in those cases (see Table S10 instead).

| **Instructed Finger** | **Ring Flexion Fatigue** | | | | | **Ring Extension Fatigue** | | | | |
|---|---|---|---|---|---|---|---|---|---|---|
| Measured Finger | Pre | Post | t | p | Estimate [95% CI] | Pre | Post | t | p | Estimate [95% CI] |
| **Index** | | | | | | | | | | |
| Index | 173.0±12.8 | 172.1±11.2 | | | | 172.2±12.2 | 171.2±9.7 | | | |
| Middle | 68.8±17.8 | 83.3±23.2 | 3.4 | 0.003 | 14.3 [5.5, 23.2] | 72.4±20.6 | 79.2±24.7 | 1.7 | 0.100 | 6.8 [-1.4, 15.1] |
| Ring | 42.4±13.1 | 59.9±27.0 | 3.8 | 0.001 | 17.6 [7.8, 27.3] | 48.4±20.9 | 52.9±24.1 | 1.3 | 0.194 | 4.5 [-2.5, 11.6] |
| Little | 34.8±8.0 | 44.8±21.0 | 2.6 | 0.018 | 10.3 [2.0, 18.6] | 38.1±14.9 | 39.5±20.5 | 0.4 | 0.671 | 1.3 [-5.2, 7.9] |
| **Middle** | | | | | | | | | | |
| Index | 59.1±25.3 | 65.9±34.0 | 1.1 | 0.271 | 6.8 [-5.8, 19.4] | 53.2±21.4 | 59.3±25.1 | 2.6 | 0.019 | 6.1 [1.1, 11.1] |
| Middle | 178.8±10.9 | 179.9±9.2 | | | | 180.1±7.5 | 181.8±7.7 | | | |
| Ring | 100.5±27.6 | 109.9±32.3 | 2.7 | 0.014 | 9.4 [2.1, 16.6] | 102.5±25.8 | 106.0±30.1 | 0.9 | 0.357 | 3.5 [-4.3, 11.4] |
| Little | 37.4±19.3 | 48.0±32.3 | 1.9 | 0.077 | 10.6 [-1.3, 22.6] | 40.1±20.8 | 42.4±22.2 | 0.6 | 0.530 | 2.3 [-5.2, 9.8] |
| **Ring** | | | | | | | | | | |
| Index | 38.5±12.5 | 40.8±16.5 | 1.1 | 0.270 | 2.3 [-1.9, 6.5] | 39.5±15.9 | 41.7±16.9 | 1.2 | 0.251 | 2.2 [-1.7, 6.1] |
| Middle | 75.1±25.1 | 95.2±38.5 | 3.0 | 0.008 | 20.1 [5.9, 34.2] | 80.6±31.2 | 89.6±34.7 | 1.3 | 0.211 | 9.0 [-5.5, 23.5] |
| Ring | 183.2±13.2 | 181.8±8.0 | | | | 186.7±9.5 | 186.1±9.8 | | | |
| Little | 55.5±27.8 | 87.4±43.3 | 4.9 | < 0.001 | 29.4 [16.7, 42.0] | 62.3±26.5 | 67.3±31.4 | 0.9 | 0.359 | 5.0 [-6.2, 16.2] |
| **Little** | | | | | | | | | | |
| Index | 38.5±11.8 | 45.0±23.2 | 1.5 | 0.159 | 6.1 [-2.6, 14.8] | 42.5±13.5 | 44.8±16.7 | 1.4 | 0.175 | 3.1 [-1.5, 7.8] |
| Middle | 52.8±28.4 | 66.9±36.5 | 2.2 | 0.045 | 11.9 [0.3, 23.5] | 59.2±31.3 | 67.4±35.8 | 2.3 | 0.031 | 9.3 [1.0, 17.6] |
| Ring | 128.0±53.9 | 136.2±48.5 | 1.6 | 0.123 | 6.3 [-1.9, 14.6] | 123.2±51.1 | 138.8±45.6 | 3.0 | 0.008 | 14.4 [4.2, 24.5] |
| Little | 175.4±14.0 | 177.2±11.5 | | | | 177.0±13.1 | 177.7±11.0 | | | |

**Table S9** Summary data (mean ± standard deviation) and pairwise comparisons pre- vs. post-fatigue (t-statistics, p-values, and estimated difference [95% confidence interval]) of peak flexion angle (°) across both visits during finger flexion-extension movements at 1 Hz. Estimates are based on linear mixed-effects model outputs and are expressed as post-fatigue relative to pre-fatigue. Please note that separate statistical tests were performed on the angles of the *instructed* fingers, hence no model outputs are presented in those cases (see Table S10 instead).

| **Instructed Finger** | **Ring Flexion Fatigue** | | | | | **Ring Extension Fatigue** | | | | |
|---|---|---|---|---|---|---|---|---|---|---|
| Measured Finger | Pre | Post | t | p | Estimate [95% CI] | Pre | Post | t | p | Estimate [95% CI] |
| **Index** | | | | | | | | | | |
| Index | 175.3±15.2 | 172.8±10.6 | | | | 171.2±13.7 | 175.4±7.9 | | | |
| Middle | 68.8±17.4 | 77.1±26.2 | 2.4 | 0.027 | 8.3 [1.0, 15.5] | 74.8±16.5 | 79.5±21.6 | 1.5 | 0.162 | 4.7 [-2.1, 11.5] |
| Ring | 42.7±14.5 | 52.2±26.9 | 2.3 | 0.032 | 9.8 [0.9, 18.7] | 47.5±16.1 | 55.8±24.7 | 2.6 | 0.019 | 8.4 [1.5, 15.2] |
| Little | 34.0±8.7 | 39.8±19.2 | 1.7 | 0.105 | 6.1 [-1.4, 13.5] | 36.0±13.8 | 42.6±20.6 | 2.9 | 0.009 | 6.5 [1.8, 11.2] |
| **Middle** | | | | | | | | | | |
| Index | 55.2±22.0 | 65.2±26.3 | 1.6 | 0.125 | 9.6 [-2.9, 22.2] | 55.0±23.3 | 62.6±22.1 | 2.9 | 0.008 | 7.6 [2.2, 13.1] |
| Middle | 179.2±11.6 | 179.4±8.5 | | | | 178.8±9.5 | 180.6±8.7 | | | |
| Ring | 99.5±26.1 | 112.4±31.7 | 2.0 | 0.055 | 12.4 [-0.3, 25.2] | 103.1±27.5 | 110.9±29.1 | 3.5 | 0.003 | 7.1 [2.8, 11.4] |
| Little | 32.8±17.6 | 48.6±29.2 | 3.0 | 0.008 | 15.6 [4.5, 26.7] | 38.2±21.4 | 41.9±19.3 | 1.5 | 0.152 | 3.7 [-1.5, 9.0] |
| **Ring** | | | | | | | | | | |
| Index | 38.1±13.3 | 41.2±18.2 | 1.0 | 0.324 | 2.9 [-3.1, 8.9] | 38.7±14.2 | 43.4±19.1 | 2.2 | 0.041 | 4.7 [0.2, 9.1] |
| Middle | 75.1±28.3 | 89.7±32.1 | 1.8 | 0.095 | 15.1 [-2.9, 33.1] | 84.7±35.2 | 93.0±41.7 | 1.3 | 0.200 | 8.4 [-4.8, 21.6] |
| Ring | 183.8±11.9 | 180.5±8.8 | | | | 186.0±7.5 | 186.5±8.9 | | | |
| Little | 51.7±15.9 | 85.3±42.2 | 4.1 | 0.001 | 31.0 [14.9, 47.1] | 58.6±30.9 | 75.4±37.9 | 3.3 | 0.003 | 16.7 [6.2, 27.2] |
| **Little** | | | | | | | | | | |
| Index | 42.7±15.2 | 44.0±19.8 | 1.0 | 0.312 | 2.8 [-2.9, 8.4] | 44.5±12.0 | 46.7±17.8 | 1.1 | 0.296 | 2.2 [-2.1, 6.5] |
| Middle | 60.6±32.1 | 64.1±31.5 | 0.9 | 0.400 | 3.0 [-4.3, 10.4] | 59.8±30.6 | 62.5±29.6 | 0.9 | 0.389 | 2.7 [-3.7, 9.1] |
| Ring | 124.8±48.1 | 130.7±49.3 | 1.6 | 0.138 | 6.8 [-2.4, 15.9] | 131.2±52.9 | 136.5±43.6 | 1.3 | 0.206 | 5.3 [-3.2, 13.7] |
| Little | 174.7±15.7 | 177.9±10.2 | | | | 175.6±13.8 | 178.0±13.9 | | | |

**Table S10:** Statistical results (t-statistics, degrees of freedom, p-values, raw estimates [90% confidence interval], and Hedge's g effect size [90% confidence interval]) from the null hypothesis significance tests and equivalence tests comparing peak flexion angle (°) by the *instructed* finger pre- vs. post-fatigue. The effect sizes are expressed as post-fatigue relative to pre-fatigue). The a priori effect size for the equivalence tests was set at 10°. Together, a non-significant null hypothesis test and a significant equivalence test allows us to conclude that the pre- and post-fatigue force magnitudes are equivalent (mean difference within ±10°).

| | **Null hypothesis significance test** | | | **Equivalence test** | | | **Effect sizes** | |
|---|---|---|---|---|---|---|---|---|
| | t | df | p | t | df | p | Raw Estimate [90% CI] | Hedge's g [90% CI] |
| ***Ring Flexion Fatigue*** | | | | | | | | |
| **0.50 Hz** | | | | | | | | |
| Index | -1.0 | 19 | 0.316 | 2.1 | 19 | 0.025 | -3.3 [-8.9, 2.2] | -0.2 [-0.6, 0.1] |
| Middle | -0.3 | 18 | 0.767 | 5.3 | 18 | < 0.001 | -0.5 [-3.6, 2.6] | -0.1 [-0.4, 0.3] |
| Ring | -0.4 | 19 | 0.679 | 4.4 | 19 | < 0.001 | -0.9 [-4.5, 2.7] | -0.1 [-0.4, 0.3] |
| Little | 0.0 | 19 | 0.971 | -5.3 | 19 | < 0.001 | 0.1 [-3.2, 3.3] | 0.0 [-0.3, 0.4] |
| **0.75 Hz** | | | | | | | | |
| Index | -0.6 | 18 | 0.542 | 4.0 | 18 | < 0.001 | -1.3 [-5.1, 2.4] | -0.1 [-0.5, 0.2] |
| Middle | 0.5 | 19 | 0.621 | -4.4 | 19 | < 0.001 | 1.0 [-2.5, 4.6] | 0.1 [-0.2, 0.5] |
| Ring | -0.6 | 19 | 0.545 | 3.7 | 19 | 0.001 | -1.4 [-5.4, 2.6] | -0.1 [-0.5, 0.2] |
| Little | 0.8 | 19 | 0.443 | -3.5 | 19 | 0.001 | 1.8 [-2.2, 5.9] | 0.2 [-0.2, 0.5] |
| **1 Hz** | | | | | | | | |
| Index | -1.1 | 19 | 0.270 | 3.4 | 19 | 0.001 | -2.5 [-6.3, 1.3] | -0.2 [-0.6, 0.1] |
| Middle | -0.2 | 18 | 0.838 | 5.4 | 18 | < 0.001 | -0.4 [-3.5, 2.7] | 0.0 [-0.4, 0.3] |
| Ring | -4.0 | 18 | 0.001 | 3.2 | 18 | 0.002 | -5.5 [-7.9, -3.2] | -0.9 [-1.3, -0.4] |
| Little | 0.4 | 18 | 0.663 | -3.4 | 18 | 0.001 | 1.1 [-3.3, 5.6] | 0.1 [-0.3, 0.5] |
| ***Ring Extension Fatigue*** | | | | | | | | |
| **0.50 Hz** | | | | | | | | |
| Index | 1.3 | 19 | 0.211 | -3.0 | 19 | 0.004 | 3.0 [-1.0, 7.1] | 0.3 [-0.1, 0.6] |
| Middle | 1.2 | 19 | 0.248 | -4.1 | 19 | < 0.001 | 2.2 [-1.0, 5.5] | 0.3 [-0.1, 0.6] |
| Ring | 0.4 | 19 | 0.698 | -6.0 | 19 | < 0.001 | 0.6 [-2.1, 3.3] | 0.1 [-0.3, 0.4] |
| Little | 0.6 | 19 | 0.540 | -3.8 | 19 | 0.001 | 1.4 [-2.5, 5.3] | 0.1 [-0.2, 0.5] |
| **0.75 Hz** | | | | | | | | |
| Index | -0.5 | 19 | 0.650 | 3.9 | 19 | < 0.001 | -1.1 [-5.0, 2.9] | -0.1 [-0.5, 0.3] |
| Middle | 1.1 | 19 | 0.267 | -5.7 | 19 | < 0.001 | 1.7 [-0.9, 4.2] | 0.2 [-0.1, 0.6] |
| Ring | -0.3 | 19 | 0.778 | 4.3 | 19 | < 0.001 | -0.6 [-4.4, 3.1] | -0.1 [-0.4, 0.3] |
| Little | 0.6 | 18 | 0.547 | -3.9 | 18 | < 0.001 | 1.3 [-2.5, 5.2] | 0.1 [-0.2, 0.5] |
| **1 Hz** | | | | | | | | |
| Index | 1.5 | 19 | 0.141 | -2.2 | 19 | 0.022 | 4.2 [-0.5, 8.8] | 0.3 [0.0, 0.7] |
| Middle | 0.9 | 19 | 0.354 | -4.1 | 19 | < 0.001 | 1.9 [-1.5, 5.3] | 0.2 [-0.2, 0.6] |
| Ring | 0.2 | 19 | 0.810 | -4.8 | 19 | < 0.001 | 0.5 [-2.9, 3.9] | 0.1 [-0.3, 0.4] |
| Little | 1.1 | 19 | 0.271 | -3.6 | 19 | 0.001 | 2.4 [-1.3, 6.1] | 0.2 [-0.1, 0.6] |
